# Unaware Processing of Words Activates Experience-Derived Information in Conceptual-Semantic Brain Networks

**DOI:** 10.1101/2024.07.15.603501

**Authors:** Marta Ghio, Barbara Cassone, Marco Tettamanti

**Affiliations:** Faculty of Mathematics and Natural Sciences, Heinrich Heine University Düsseldorf, Düsseldorf, Germany; CIMeC - Center for Mind/Brain Sciences, University of Trento, Italy; Department of Psychology, University of Milan-Bicocca, Milano, Italy

**Keywords:** Conceptual-semantic representations, experience, awareness, continuous flash suppression, manipulable objects, emotions

## Abstract

The recognition of manipulable objects results from the encoding of sensory input in combination with predictive decoding of experience-derived visuomotor information stored in conceptual-semantic representations. This grounded interpretive processing was previously found to subsist even under unaware perception of manipulable object pictures. In this fMRI study, we first aimed to extend this finding by testing whether experientially grounded visuomotor representations are unawarely recruited when manipulable objects are not visually depicted, but only referred to by words presented subliminally through continuous flash suppression. Second, we assessed the generalizability of decoding experience-derived conceptual information to other semantic categories, by extending our investigation to subliminally presented emotion words and testing for unaware recruitment of grounded emotion representations in the limbic system. Univariate analysis of data sampled from 21 human participants (14 females) showed that manipulable object words selectively activated a left-lateralized visuomotor network, both when words were presented below perceptual threshold and when participants subjectively reported lack of stimulus awareness. Emotion words selectively engaged the bilateral limbic network, although univariate analysis did not provide evidence for its recruitment under subliminal perceptual conditions. In turn, multivariate pattern analysis showed that neural codes associated with both manipulable object and emotion words could be decoded even in the absence of perceptual awareness. These findings suggest that the brain automatically engages in conceptual-semantic decoding of experience-derived information not only when circumstances require to interact with manipulable objects and emotions, but also when these referents are dislocated in time and space and only referred to by words.

## 1. Introduction

Our brain constantly receives sensory input from around and within us and reacts to it, also in anticipatory mode, by activating conceptual-semantic representations derived from previous experience with similar percepts. For example, visual perception of manipulable objects (MO) engages not only bottom-up visual encoding but also top-down conceptual-semantic decoding based on stored MO concepts (Noppeney et al., 2006). These are considered to encompass high-level semantic (e.g., categorical) as well as visual, motor, and functional information derived from our previous experience with MO exemplars (Binder & Desai, 2011; Lambon Ralph et al., 2017). The representation, storage and retrieval of visual properties rely on bilateral ventral occipito-temporal areas, while motor and functional properties draw on portions of the left-hemispheric parietal and premotor cortices that support the motor programming of actions related to the object function (Ishibashi et al., 2016; Chen et al., 2017). This MO-specific visuomotor network (MOVN) interacts with the general semantic brain network (Binder et al., 2009) to form inferences about the incoming visual input that allow us to ultimately identify the object and possibly to plan a motor response (Binder & Desai, 2011; Lambon Ralph et al., 2017).

Notably, MO conceptual-semantic representations seem to be activated also under unaware visual perception conditions (Ludwig et al., 2016; Fahrenfort et al., 2017) such as those instantiated by Continuous Flash Suppression (CFS, Tsuchiya & Koch, 2005). By applying CFS in combination with fMRI, we showed the recruitment and effective connectivity of the MOVN both under aware and unaware processing of MO pictures (Tettamanti et al., 2017; Ghio et al., 2022). A possible concern, however, is that the MOVN was recruited due to the visual processing of irrelevant perceptual properties such as the prototypical elongated shape of MO, rather than to the retrieval of semantic features such as manipulability and affordance. The first aim of the current study was to address this concern by evaluating the subliminally-induced activation of the MOVN in processing MO stimuli that lack a visual depiction of the objects, as is the case for words. While both pictorial and verbal stimuli trigger conceptual-semantic processing, the latter are arbitrarily associated with their referents through learning and do not carry any object-related visuomotor information. A recruitment of the MOVN under subliminal processing of MO words would thus speak for the hypothesis that conceptual-semantic processing activating experience-derived MO information occurs automatically.

The second aim was to examine the generalizability of the hypothesis that experience-derived information constituting conceptual-semantic representations is automatically recruited in subliminal perception, by extending our investigation to emotions (EM). Conceptual representations of discrete EM (e.g., *happiness, anger*) are characterized by valence, arousal, interoceptive and somatic information derived from our previous experience of EM instances (Ghio & Tettamanti, 2016; Borghi et al., 2018; Conca et al., 2021). This information is supposed to be represented and stored in the same limbic network that supports the experience of EM and comprises, bilaterally, the amygdala, insula, anterior cingulate, and orbitofrontal cortices, putamen, and midbrain (Lindquist et al., 2012). This EM-specific limbic network (EMLN) has been shown to be activated when we visually perceive an instance of an EM in, e.g., a face picture, even presented below the perceptual threshold (Jiang & He, 2006; Lindquist et al., 2012; Dahlén et al., 2022). As for MO, however, to overcome the potential criticism that subliminal activation is due to visual properties of the pictures, we used subliminally presented EM words. Previous behavioural and ERP studies using CFS proved that EM word meaning is also accessed under subliminal conditions (Lei et al., 2017; Sklar et al., 2012; Vinson et al., 2011; Yang & Yeh, 2011), without providing any insight, however, into the automatic recruitment of the EMLN.

In the present fMRI study, we tested the aforementioned hypothesis and its generalization with CFS of MO and EM words. Univariate and multivariate pattern analyses showed automatic activation of MOVN and EMLN during, respectively, MO and EM word processing.

## 2. Methods

### 2.1. Experimental design and hypotheses

This fMRI study aimed at assessing the hypothesis that experience-derived information constituting MO (e.g., *hammer*) and EM (e.g., *envy*, *joy*) conceptual-semantic representations is automatically recruited in subliminal perception. To this purpose we tested whether the subliminal processing of MO and EM words specifically engages, respectively, the MOVN and the EMLN (see Fig. 1A and the “Data analysis” section below for the definition of the specific regions of interest and coordinates). Because valence is a dominant feature for EM conceptual-semantic representations (Conca et al., 2021), with respect to which EM words are consistently characterized across languages (Jackson et al., 2019), EM words were parametrically distributed along the negative-to-positive valence dimension, including an equal number of negative (EMneg) and positive (EMpos) words. In addition, neutral words NEU (e.g., *thought*) served as a control condition to test the effect of valence within the EMLN.

**Figure 1.**
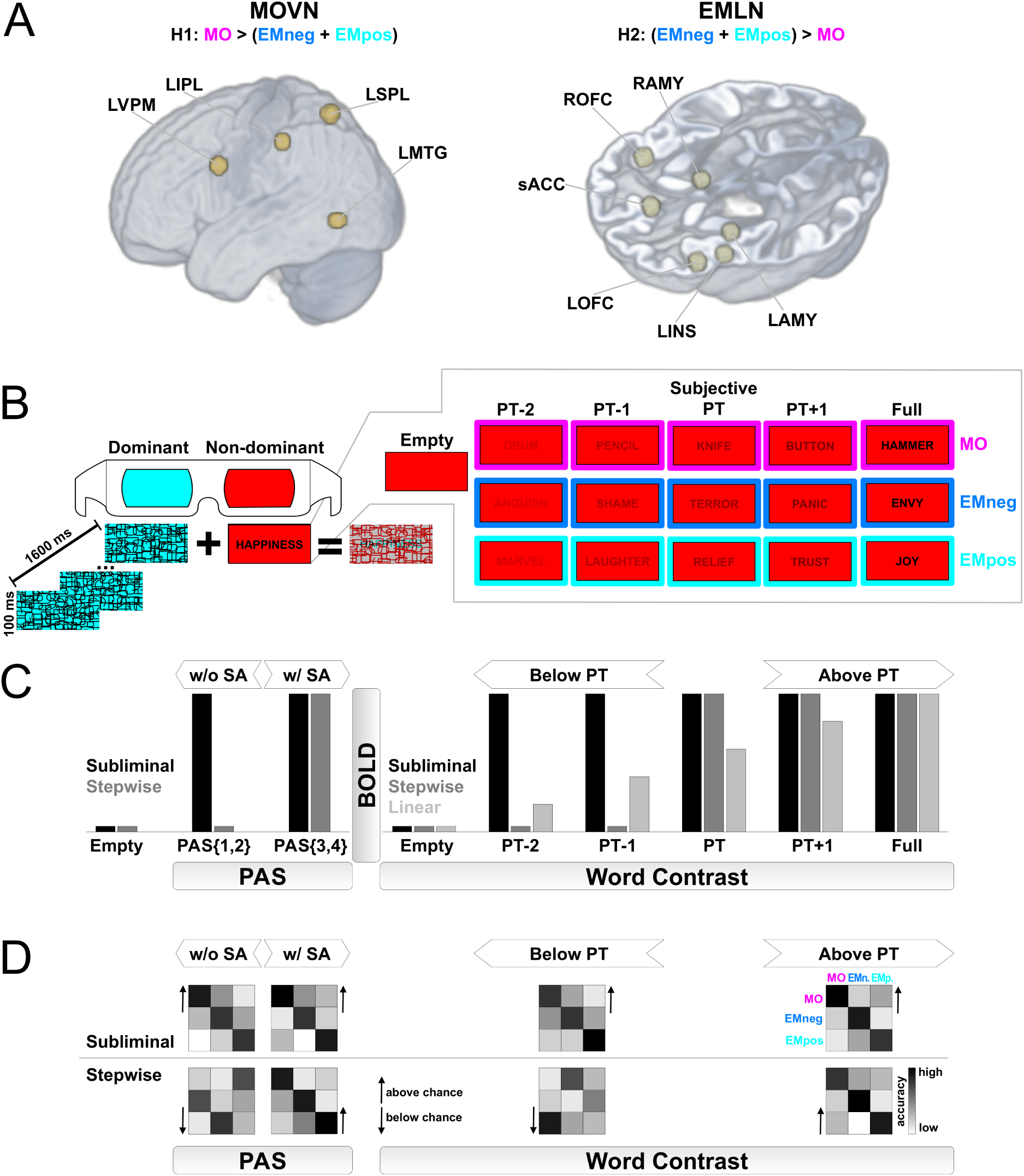
Experimental hypotheses and procedures. A) We tested the hypothesis (H1) that, in comparison to emotion (EM) words with either a negative (EMneg) or a positive (EMpos) valence, subliminally perceived manipulable object (MO) words selectively activated the MO-specific visuomotor network (MOVN), constituted by the left ventral premotor cortex (LVPM), the left inferior parietal lobule (LIPL) the left superior parietal lobule (LSPL), and the left middle temporal gyrus (LMTG). Conversely, our second hypothesis (H2) predicted that stronger activations for the subliminal perception of EMneg and EMpos compared to MO would be found in the EM-specific limbic network (EMLN), comprising the left amygdala (LAMY), the right amygdala (RAMY), the left orbito-frontal cortex (LOFC), the right orbito-frontal cortex (ROFC), the left anterior insula (LINS), and the sub-/pre-genual anterior cingulate cortex (sACC). For both the H1 and H2 hypotheses, the alternative hypothesis (H0) was that a lack of selective activation in the MOVN or EMLN networks for subliminal word processing would undermine the view that conceptual semantic representations encompass experience-derived information. Anatomical spheres of interest of 6 mm radius are rendered on a volumetric average brain image of all participants. B) Experimental trials consisted in the presentation of a red-filtered written word (word types: MO, EMneg, EMpos), which was only visible through the red anaglyph lense positioned on the non-dominant eye. The word was presented either with null visual contrast (Empty baseline), at subjective perceptual threshold (PT), and below (PT-2 and PT-1) or above (PT+1, Full) that threshold. These subliminal or supraliminal visual contrast levels resulted from Continuous Flash Suppression, instantiated through cyan-filtered mask overlays flickered at 10 Hz, which were only visible through the cyan anaglyph lense positioned on the dominant eye. C) fMRI data were inspected through univariate analysis for the presence of, respectively, MO-specific BOLD responses in the MOVN or EM-specific responses in the EMLN, fitting a “subliminal”, as opposed to a “stepwise” or “linear”, profile of activation as a function of Word Contrast (i.e., below or above PT), and as a function of PAS rating (i.e., with or without subjective awareness (SA)). D) Multivariate analysis of brain activation patterns sought for accurate (i.e., above chance) decoding of Word Types, following a “subliminal” as opposed to a “stepwise” decoding profile; that is, a successful decoding both below and above PT, and both with and without SA.

To instantiate subliminal word processing we applied a CFS paradigm (Fig. 1B). After determining the perceptual threshold for each participant, the word stimuli (Word Type: MO, EMneg, EMpos, NEU) were presented at five incremental visual contrast levels (Word Contrast: two levels below, one at participant’s perceptual threshold, and two above), with void stimuli as perceptual control. Our CFS paradigm also required to indicate subjective ratings along a perceptual awareness scale (PAS; Ramsøy and Overgaard, 2004), which allowed us to determine the presence or absence of subjective perceptual awareness, independently of word contrast.

We first applied univariate analyses (Fig. 1C) to assess subliminal fMRI activation (1) as a function of Word Contrast (reflecting the subliminal vs. supraliminal conditions defined based on participant’s perceptual threshold), and (2) as a function of PAS ratings (reflecting the subjective perceptual awareness). In both analyses we tested the hypothesis that the MOVN and the EMLN specifically respond to, respectively, MO and EM words following a “subliminal” activation amplitude profile, which was defined as stable activation either (1) across the five incremental contrast levels or (2) with and without subjective awareness. The subliminal profile, which would speak for unaware processing, was compared against a “linear” profile, i.e., activation increase with increasing word contrast (only in 1) and a “stepwise” profile, i.e., drop of activation either (1) for word contrast below perceptual threshold or (2) without subjective awareness. In addition, for EM we also tested the effects of valence by distinguishing EMneg, EMpos and NEU.

Secondly, we applied multivariate pattern analyses (Fig. 1D), again (1) as a function of Word Contrast and (2) as a function of PAS ratings. We hypothesized that the involvement of MOVN and EMLN for the representation and subliminal processing of, respectively, MO and EM words would be reflected by the successful decoding of the word types (1) not only above but also below perceptual threshold and (2) not only with but also without subjective perceptual awareness.

### 2.2. Participants

Twenty-five Italian native speakers voluntarily participated in the study. Two participants were excluded due to the finding of brain structural anomalies. Two other participants did not comply with the CFS task and were discarded from the analyses. The sample of the remaining 21 subjects comprised 7 male and 14 female participants (mean age = 22.31 years, SD = 4.01; mean years of education = 15.76, SD = 3.25). All participants reported to be right-handed, which was in line with the Edinburgh inventory assessment (Oldfield, 1971; mean score = 0.80, SD = 0.16, with 20 participants scoring in the range from 0.5 to 1 and thus classified as right-handed and one participant scoring 0.4 and classified as ambidextrous). The Miles test (Miles, 1929) was administered for evaluating eye dominance for the CFS task: 8 participants were left, and 13 right eye dominant. All participants reported neither sensory deficits nor developmental, neurological, or psychiatric diagnoses. No participants had reading deficits according to their performance with real word (mean score = 100%, SD = 0%) and pseudoword (mean score = 99%, SD = 1%) reading tests included in the Italian battery for assessing aphasic deficits (Miceli, 1994). All participants gave written consent and received a reimbursement of 30 euro for their participation in the study. The study was approved by the ethics committee of the University of Trento, Italy.

### 2.3. Stimuli

#### 2.3.1. Word stimuli for the fMRI CFS task

As experimental word stimuli we used 75 nouns referring to MO and 75 to EM. As described above, EM nouns were parametrically distributed along the negative-to-positive valence dimension, including 25 EMneg and 25 EMpos nouns, with additional 25 NEU nouns as control condition for testing the valence effects. EMneg and EMpos were defined as referring to “a mental state that could be felt” (e.g., *anger* or *joy*; Jackson et al., 2019), also known as emotion-label nouns (Conca et al., 2021). Emotion-laden nouns (e.g., *war* or *party*) were avoided. NEU were defined as referring to “neutral mental states” (Jackson et al., 2019) that ”can not be felt” (e.g., *thought*). In line with these definitions, we checked that in the lexical database Multiwordnet (Pianta et al., 2002) all EMneg, EMpos, and NEU nouns are classified as psychological features or factotum (semantic domain level), but only EMneg and EMpos are defined as emotions or traits/attitudes (affective level). The complete list of stimuli is available as Supplementary Table 1A.

To validate the variability of EM words along the negative-to-positive valence dimension, all nouns were rated on a bipolar Likert scale (-3 = negative, 0 = neutral, +3 = positive) by an independent group of 16 volunteers (10 female and 5 male participants, 1 did not specify the sex; mean age = 29.25 years, SD = 5.48). Results showed that EMneg, EMpos, and NEU nouns were clearly distributed along the negative-to-positive valence dimension, and significantly differed from MO (Table 1A for summary statistics and Supplementary Table 1B for single word rating scores). Ratings for arousal (1 = calm or relaxation, 7 = high excitement or agitation) were also collected, as valence and arousal are considered to be independent dimensions of the semantic space (Osgood & Suci, 1955). Results showed that EMpos, NEU, and MO did not differ in arousal, although all were lower that EMneg (Table 1A). Finally, ratings on a unipolar Likert scale for imageability (1 = barely imaginable, 7 = easily imaginable), which is considered as a proxy for concreteness (Hoffman, 2016), showed that EM and NEU were less imageable than MO words. Experimental stimuli were rated for valence, arousal and imageability also by all (but one) fMRI participants (on average 8.55 days, SD = 9.97, after the MRI session). The results showed a high correlation between the ratings of the fMRI and independent participant samples (Spearman’s correlation coefficient: valence: rho = 0.971; arousal: rho = 0.937; imageability: rho = 0.994, all p values < .001).

**Table 1.**
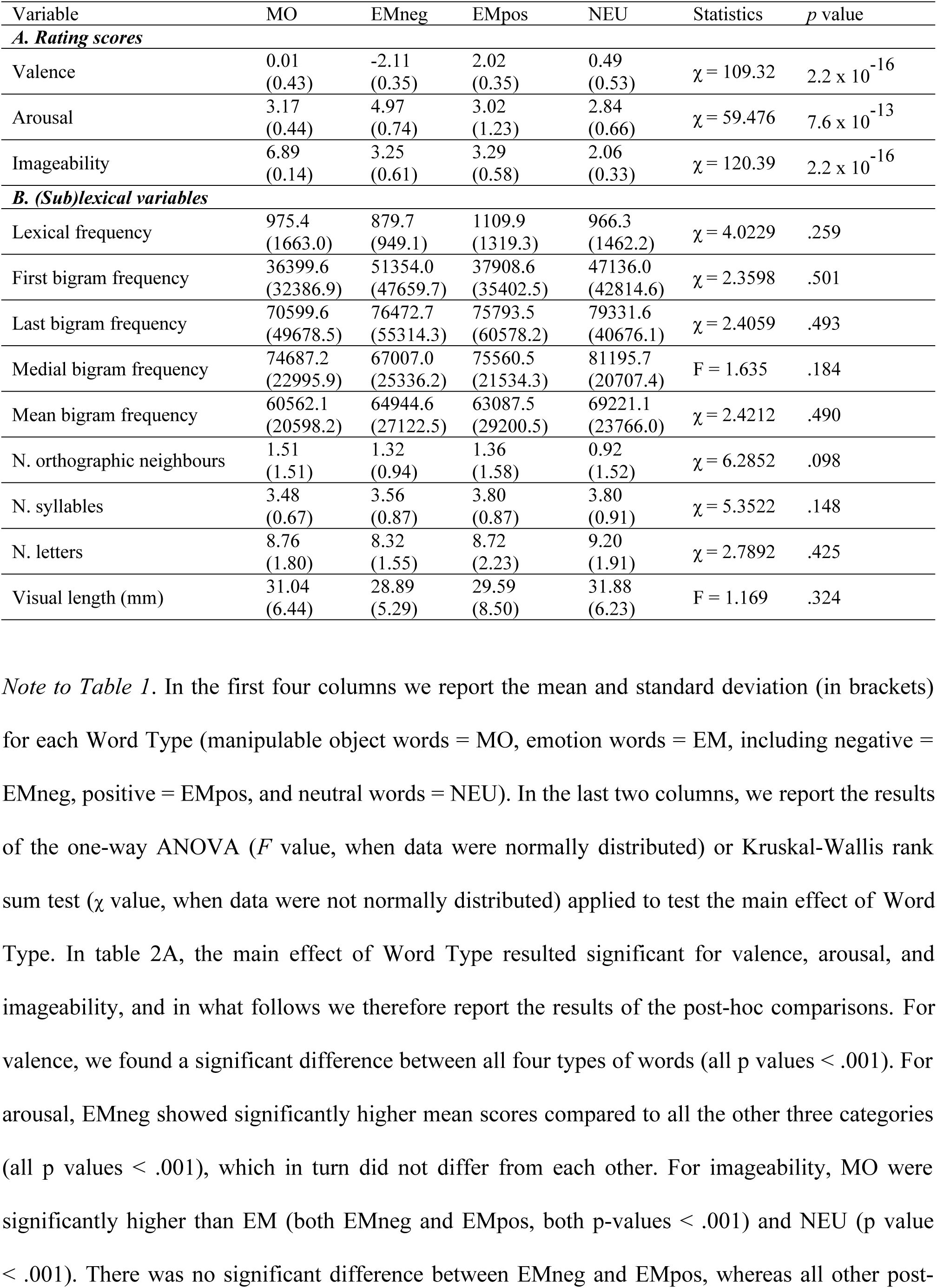
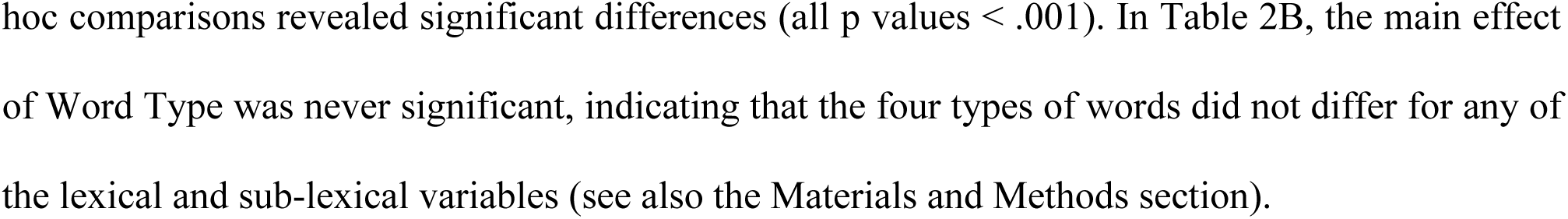
Descriptive and inferential statistics for rating scores and (sub)lexical variables characterizing the experimental stimuli.

| Variable | MO | EMneg | EMpos | NEU | Statistics | *p* value |
| --- | --- | --- | --- | --- | --- | --- |
| **A. Rating scores** |  |  |  |  |  |  |
| Valence | 0.01<br>(0.43) | -2.11<br>(0.35) | 2.02<br>(0.35) | 0.49<br>(0.53) | $\chi = 109.32$ | $2.2 \times 10^{-16}$ |
| Arousal | 3.17<br>(0.44) | 4.97<br>(0.74) | 3.02<br>(1.23) | 2.84<br>(0.66) | $\chi = 59.476$ | $7.6 \times 10^{-13}$ |
| Imageability | 6.89<br>(0.14) | 3.25<br>(0.61) | 3.29<br>(0.58) | 2.06<br>(0.33) | $\chi = 120.39$ | $2.2 \times 10^{-16}$ |
| **B. (Sub)lexical variables** |  |  |  |  |  |  |
| Lexical frequency | 975.4<br>(1663.0) | 879.7<br>(949.1) | 1109.9<br>(1319.3) | 966.3<br>(1462.2) | $\chi = 4.0229$ | .259 |
| First bigram frequency | 36399.6<br>(32386.9) | 51354.0<br>(47659.7) | 37908.6<br>(35402.5) | 47136.0<br>(42814.6) | $\chi = 2.3598$ | .501 |
| Last bigram frequency | 70599.6<br>(49678.5) | 76472.7<br>(55314.3) | 75793.5<br>(60578.2) | 79331.6<br>(40676.1) | $\chi = 2.4059$ | .493 |
| Medial bigram frequency | 74687.2<br>(22995.9) | 67007.0<br>(25336.2) | 75560.5<br>(21534.3) | 81195.7<br>(20707.4) | $F = 1.635$ | .184 |
| Mean bigram frequency | 60562.1<br>(20598.2) | 64944.6<br>(27122.5) | 63087.5<br>(29200.5) | 69221.1<br>(23766.0) | $\chi = 2.4212$ | .490 |
| N. orthographic neighbours | 1.51<br>(1.51) | 1.32<br>(0.94) | 1.36<br>(1.58) | 0.92<br>(1.52) | $\chi = 6.2852$ | .098 |
| N. syllables | 3.48<br>(0.67) | 3.56<br>(0.87) | 3.80<br>(0.87) | 3.80<br>(0.91) | $\chi = 5.3522$ | .148 |
| N. letters | 8.76<br>(1.80) | 8.32<br>(1.55) | 8.72<br>(2.23) | 9.20<br>(1.91) | $\chi = 2.7892$ | .425 |
| Visual length (mm) | 31.04<br>(6.44) | 28.89<br>(5.29) | 29.59<br>(8.50) | 31.88<br>(6.23) | $F = 1.169$ | .324 |

*Note to Table 1.* In the first four columns we report the mean and standard deviation (in brackets) for each Word Type (manipulable object words = MO, emotion words = EM, including negative = EMneg, positive = EMpos, and neutral words = NEU). In the last two columns, we report the results of the one-way ANOVA ( $F$ value, when data were normally distributed) or Kruskal-Wallis rank sum test ( $\chi$ value, when data were not normally distributed) applied to test the main effect of Word Type. In table 2A, the main effect of Word Type resulted significant for valence, arousal, and imageability, and in what follows we therefore report the results of the post-hoc comparisons. For valence, we found a significant difference between all four types of words (all $p$ values $< .001$ ). For arousal, EMneg showed significantly higher mean scores compared to all the other three categories (all $p$ values $< .001$ ), which in turn did not differ from each other. For imageability, MO were significantly higher than EM (both EMneg and EMpos, both $p$ -values $< .001$ ) and NEU ( $p$ value $< .001$ ). There was no significant difference between EMneg and EMpos, whereas all other post-

Experimental stimuli were balanced for relevant sub-lexical and lexical variables (Table 1B for detailed descriptive and inferential statistics). Lexical frequency, which has been demonstrated to influence speed in word recognition (Brysbaert et al., 2016; but see Heyman & Moors, 2014), was extracted from the SUBTLEX-IT database (Crepaldi et al., 2015) and matched across the four types of words. Moreover, since sub-lexical processing has also been found to affect word recognition (Westbury & Buchanan, 2002), MO, EMneg, EMpos, and NEU were matched for the frequency of the first, last, and average medial bigrams of each word, plus the ove rall average bigram frequency. Bigram frequencies were extracted from the PhonItalia database (Goslin et al., 2014). Stimuli were also matched for their number of orthographic neighbours (again extracted from the PhonItalia database, Goslin et al., 2014), for it has been demonstrated that a large orthographic neighbourhood size makes words easier to read (Duñabeitia et al., 2008). Finally, stimuli were balanced for their number of syllables, number of letters, and visual length calculated in millimeters using the software GIMP 2.10.4 (www.gimp.org).

#### 2.3.2. Mask stimuli for the CFS task

We used MATLAB R2017b to create 80 mask images with cyan background to make the mask visible only to the dominant eye when wearing the red-cyan anaglyph goggles (see below). Over the cyan background we displayed black patterns, which were similar in shape (e.g., closed unfilled geometrical shapes such as ellipses and rectangles, and open u-shaped figures with different orientations) and perceptual features (e.g., line width, dimension) to the Latin alphabet letters of the word stimuli (Kido & Makioka, 2014; Axelrod et al., 2015). Then we overlayed a black frame and a black fixation cross at the centre of every mask, using GIMP 2.10.4 (www.gimp.org).

### 2.4. Experimental procedure

Participants first performed a training session to familiarize with the CFS task outside the MRI scanner room. Then, they entered into the MRI scanner and performed a pre-fMRI behavioral session of the CFS task to assess their perceptual threshold in the same contextual and perceptual conditions as for the fMRI CFS task. Finally, they performed the experimental CFS task session, while their brain activity was measured with fMRI. In all these three sessions, participants wore MRI-compatible red-cyan anaglyph goggles, with the cyan lens placed above the dominant eye. The CFS trial structure was identical across the three sessions, as well as the use of Psychtoolbox-3 (v3.0.14, running in MATLAB 2017b) for stimulus delivery and behavioural response collection.

#### 2.4.1. CFS trial

Each CFS trial began with the 200-ms presentation of a fixation cross on a red background overlaid by an empty cyan image with 50% transparency (alpha blending). Then, a single word was presented in black on a red background to make it visible only to the non-dominant eye when wearing the red-cyan anaglyph goggles. Each word was written in uppercase (font: Helvetica, font dimension: 80) and displayed in the center of the screen at a trial-specific (see below) visual contrast level, for a duration of 1600 ms. In the same 1600-ms time interval, a set of 16 different masks presented at 10 Hz was overlaid on the word with 50% transparency (Fig. 1B). For every trial, the 16 masks were randomly extracted from the whole pool of 80 mask images and then presented in a randomized order. To reduce after image effects (Hong & Blake, 2009), which can occur at the offset of the last mask, we displayed for 1000 ms an image with multiple white capital X letters on a black background. After a variable within-trial delay (pseudo-randomized duration of either 1425, 1925, or 2425 ms in 4:2:1 proportion), participants were shown the display “1 2 3 4” (duration 1000 ms), which represented the four level of the PAS (Ramsøy & Overgaard, 2004): 1 = no experience; 2 = brief glimpse without recognition of any letter; 3 = almost clear perception, recognition of one or more letters, but not of the entire word; 4 = absolutely clear perception and recognition of the entire word. Participants were asked to press one of the four buttons of a response pad according to their subjective level of visual perception of the word. To prevent explicit semantic processing, questions assessing the objective perception of stimuli were avoided. The variable within-trial delay was introduced in order to reduce the temporal correlation between the stimulus visual presentation and the motor response to the PAS scale, thus preventing possible biases in interpreting BOLD signal fluctuations. At the end of the trial, a delay of variable duration (short: 2925 or 3425 ms, medium: 4075 or 4575 ms, long: variable from 5075 to 10575 ms, in 4:2:1 proportion) was included before the start of the next trial. Within-trial and between-trial delays were determined by OPTseq2 (surfer.nmr.mgh.harvard.edu/optseq) to maximize the sensitivity of the hemodynamic signal in the fMRI CFS event-related design (Dale, 1999). Trial timings were kept identical also for the training and the pre-fMRI behavioral sessions.

#### 2.4.2. Training session

Before entering the MRI scanner, participants were trained to perform the CFS task on a laptop computer. After performing 6 practice trials, participants started the training session, which comprised 105 consecutive trials. For 100 trials, we used nouns referring to jobs and careers (e.g., *biologist*), which were matched with the experimental stimuli with respect to lexical frequency, number of syllables, and number of letters (all *p* > .072; Supplementary Table 2A,B). These nouns were presented at a specific contrast level according to the following procedure (similarly to Tettamanti et al., 2017). For each participant, the 100 nouns were randomly ordered in a list and then sequentially drawn without replacement from such list and assigned, in batches of 5, to 1 of 20 transparency levels (alpha blending value ranging from 8 to 255 = fully opaque, by steps of 13 units). In addition, for 5 trials we applied a transparency level with alpha = 0, which resulted in 5 empty trials (i.e., trials devoid of words, only including the red background and the flickering cyan masks). The presentation order of the 105 trials was pseudo-randomized across participants.

#### 2.4.3. Pre-fMRI behavioral session: psychophysical assessment of perceptual threshold

The perceptual threshold of each participant was assessed before the fMRI data acquisition session. Participants were positioned in the MRI scanner and performed the CFS task on 105 consecutive trials, including 5 empty trials and 100 trials, during which 100 nouns referring to living/natural items (e.g., *elephant*, *sunflower*, *storm*, *forest)* were presented at 20 different alpha transparency levels (defined as in the training session described above). This set of nouns was matched with the experimental stimuli with respect to lexical frequency, number of syllables, number of letters, bigram frequency (initial, final, medial, average), and number of orthographic neighbours (all *p* > .103; Supplementary Table 3A,B).The same presentation procedure as in the training session was applied.

To estimate the subjective perceptual threshold, we adopted the same procedure as in Tettamanti et al. (2017). First, for each participant we plotted the number of responses with a PAS score equal to 3 or 4 (indicating high to maximal subjective perceptual awareness) for each of the 20 transparency levels plus the empty trials. By using nonlinear least-squares estimation in MATLAB 2017b, we fitted these values with a psychometric sigmoid function f = L + (R – L) / {1 + exp [– (x – i) / W]}, where x = transparency level, L = left horizontal asymptote (initial value = 0); R = right horizontal asymptote (initial value = 5), i = point of inflection (initial value = 2.5), and W = width of the rising interval (initial value = 1). We thus defined the subjective perceptual threshold (PT) as the first transparency level above the estimated point of inflection. Based on the subjective perceptual threshold, five transparency levels were individually tailored to be used in the fMRI session (and henceforth defined as Word Contrast levels). Specifically, two tailored contrast levels were below PT: one defined as PT -25% (PT-2) and the other as PT -10% (PT-1). One contrast level was at PT. Two tailored contrast levels were above PT: one defined as PT +10% (PT+1), while the other was set for all participants at the maximum available contrast (Full, i.e., corresponding to alpha = 255). In addition, for all participants, we included a null contrast control condition (Empty), consisting of a red square devoid of any words. For participants with a subjective PT equal or lower than the transparency alpha level value of 69, the contrast levels below PT had to be tailored to a narrower range. Specifically, for participants (n = 9) with a subjective PT between alpha = 36 and 69, we set PT-2 to alpha = 5. For participants (n = 4) with a subjective PT lower than alpha = 36, we set PT-2 to alpha = 5 and PT-1 to alpha = 10.

#### 2.4.4. fMRI CFS task

The fMRI experiment consisted in a within-subject event-related experimental design, where Word Type (MO, EMneg, EMpos, NEU) and Word Contrast (PT-2, PT-1, PT, PT+1, Full) were factorially manipulated. All participants completed 6 runs. Every run comprised 55 trials: 5 empty, 25 MO and 25 EM trials. All the 25 EM words in every run belonged to the same valence category (either EMneg, or EMpos, or NEU), and the EM category assignment to the runs was counterbalanced for every participant on the basis of a Latin square, such that all words of the three categories occurred once in the first three runs before being extracted again in the second three runs. Each of the 75 MO words were presented once in the first three runs and a second time in the last three runs. Thus, every word was presented twice throughout the experiment. Within every run, 5 MO and 5 EM were presented at one of the 5 word contrast levels. The presentation order of the different Word Type × Word Contrast combinations was unique for every participant and run, and determined by OPTseq2. The assignment of the specific word stimuli to the Word Type × Word Contrast combinations was pseudo-randomized for every participant and run, in order to minimize the number of stimuli presented at the same contrast in their first occurrence (during one of the first three runs) versus their second occurrence (during one of the last three runs) for a given participant.

### 2.5. MRI data acquisition

MRI scans were acquired with a 3 Tesla Siemens Magnetom Prisma whole body MRI scanner using a 64-channels head coil. During the PT estimation session, we obtained a T1-weighted MPRAGE anatomical scan (288 slices, TR = 3280 ms, TE = 2.01 ms, flip angle = 7°; FOV = 252 × 252 mm, slice thickness = 0.7 mm). During the experimental CFS task session, whole-brain functional images were acquired with a T2*-weighted gradient-echo, EPI pulse sequence, using BOLD contrast (TR = 1000 ms, TE = 28 ms, flip angle = 59°, multi-band acceleration factor = 5). Each functional image comprised 65 transversal slices (2 mm thick), acquired in interleaved mode (FOV = 200 x 200 mm). Each participant underwent 6 consecutive functional scanning sessions, each comprising 520 scans and lasting 8 min and 50 s.

The stimuli were projected on a computer monitor located in the back of the scanner and visible to the participants through a mirror placed on the head coil above their eyes. An MRI- compatible fibre-optic response box with four buttons allowed the participants to give PAS score responses using their left hand (index finger on the “4” button). The left hand was chosen to avoid contamination of the BOLD signal acquired in left hemisphere where language processing is lateralized, even though the overlapping between motor responses and word presentation was anyway minimized thank to the within-trial delay.

### 2.6. Data analysis

#### 2.6.1. Statistical analysis of behavioural data

To evaluate the effects of Word Type and Word Contrast on the PAS scores during fMRI, we ran a series of Generalized Linear Mixed Models using R 4.2.2 (R Core Team, 2022). The first simplest model comprised PAS scores as dependent variable, the six runs as fixed-effect confounds, and the participants as random-effect factor. Word stimuli were also modelled as random-effect factor, although we note that, since their variance equaled zero (SD = 0), their inclusion or omission from the model yields identical results (Pasch et al., 2013). We then specified a series of models with increasing complexity by adding Word Type (MO, EMneg, EMpos, NEU), Word Contrast (PT-2, PT-1, PT, PT+1, Full), and the Word Type by Word Contrast interaction as fixed-effect factors. All models were fit to a Poisson distribution, with a log-link function. To evaluate whether each additional predictor significantly increased the model fit, we ran a χ2 loglikelihood ratio test (declared significance α level: .05) between each hierarchically more complex model and the simpler one.

#### 2.6.2. Univariate statistical analyses of fMRI data

For MRI data preprocessing and univariate statistical analyses, we used SPM12 v7771 (www.fil.ion.ucl.ac.uk/spm). Structural MRI images were segmented and registered to the Montreal Neurological Institute (MNI) standard space. Functional images were corrected for geometrical distortions, spatially realigned, corrected for slice timing, and normalized to the MNI space by using the segment procedure with the subject-specific segmented structural images as customized segmentation priors. Functional images were spatially smoothed with a 4 mm FWHM Gaussian kernel.

By applying a two-stage random-effects statistical approach, we performed univariate statistical analyses of fMRI activation (1) as a function of Word Contrast and (2) as a function of PAS rating. For all the reported univariate analyses, the significance threshold was declared at peak level *p* = .05, using a small volume familywise error (FWE) type correction for multiple comparisons, with spherical small volumes of 6 mm radius centered on the coordinates of interest (x, y, z cartesian values in mm referring to the MNI space) within the MOVN and the EMLN (Fig. 1A). The coordinates for the MOVN were taken from the meta-analysis by Ishibashi et al. (2016), and included the left ventral premotor cortex (LVPM; x = -50.3, y = 6.3, z = 30.7), the left inferior (LIPL; x = -45.7, y = -31.7, z = 44.8) and superior parietal lobules (LSPL; x = -23.4, y = -61.1, z = 60.9), and the left middle temporal gyrus (LMTG; x = 17.8, y = 65.1, z = 1.8). For EMLN, the coordinates were taken from Lindquist et al. (2012), and included the left (LAMY; x = -20, y = -4, z = -16) and right amygdala (RAMY; x = 22, y = -4, z = -16), the left (LOFC; x = -40, y = 26, z = -6) and right orbito-frontal cortex (ROFC; x = 42, y = 24, z = -4), the left anterior insula (LINS; x = - 38, y = 8, z = -10) and the sub-/pre-genual anterior cingulate cortex (sACC; x = 0, y = 38, z = 6).

##### 2.6.2.1. Univariate statistical analysis as a function of Word Contrast

At the first level, for each participant we specified a general linear model, which comprised six separate runs, each including 11 regressors of interest: Empty, MOPT-2, MOPT-1, MOPT, MOPT+1, MOFull, EMPT-2, EMPT- 1, EMPT, EMPT+1, EMFull, with EM corresponding to either EMneg, EMpos, or NEU depending on the run. Evoked responses were aligned to the onset of each trial (appearance of the fixation cross), and their duration was fixed to 2800 ms (comprising 200 ms for the presentation of the fixation cross, 1600 ms for the presentation of the masked stimuli, and 1000 ms for the presentation of the anti-after effect image). Experimental confounds, namely PAS responses (for each trial, onset aligned to the display of the PAS scale and duration equal to the interval between this time point and the participant’s response), task instructions, and head movement parameters yielded by image realignment, were modelled as separate regressors. If there were trials in which participants did not answer to the PAS scale (misses) or answered before the PAS question was displayed, these were modelled in the same confound regressor with task instructions. For all experimental conditions, we modelled the evoked responses as canonical haemodynamic response functions. The time series were high-pass filtered at 128 s and pre-whitened by means of an autoregressive model AR(1). Global normalization was performed to account for possible differences in signal due to the presentation of EMneg, EMpos, and NEU stimuli in different runs. Statistical analyses were restricted by using an explicit mask including only the voxels with gray matter tissue probability > .1, as determined based on the segmented structural images of each participant.

We defined two different random-effects, full factorial designs. The first one was aimed at testing the main effect of Word Type. Accordingly, within the estimated first-level general linear model we defined a series of 20 condition-specific contrasts with the Empty baseline subtracted (e.g., MOPT-2 – Empty). Namely, we defined a weight of +1 for the regressor of the specific condition of interest (e.g., MOPT-2), a weight of -1 for the Empty baseline and a weight of 0 for all other regressors. At the second-level, we entered the resulting 20 first-level contrasts into a model including Word Type (4 levels: MO, EMneg, EMpos, NEU) and Word Contrast (5 levels: PT-2, PT-1, PT, PT+1, Full) as within-subject factors. We assumed dependence and equal variance between the levels of both factors. Within this full factorial design, we specified a series of T- contrasts. Specifically, to examine the effects of MO we tested (i) MO – (EMneg+EMpos), whereas to examine the effects of EM, we tested: (ii) (EMneg+EMpos) – MO. In addition for EM we tested (iii) the simple effect of valence (i.e., with linear contrast weights: -1 for EMneg, 0 for NEU, and +1 for EMpos); (iv) the quadratic effect of valence (i.e., with non-linear weights: +1 for EMneg, -2 for NEU, and +1 for EMpos).

The second random-effects, full factorial design aimed at testing whether the effects of MO and EM as a function of Word Contrast followed either a subliminal (i.e., activation both above an d below perceptual threshold), a linear (i.e., gradual activation increase with contrast), or a stepwise (i.e., activation only above perceptual threshold) profile (Fig. 1C). In order to model these three different profiles, within the estimated first-level General Linear Model, we defined 24 condition- specific contrasts with the Empty baseline not subtracted but explicitly modeled. Namely, we defined a weight of +1 for the regressor of the specific condition of interest (e.g., MOPT-2) and a weight of 0 for all the other regressors. At the second level, we entered the resulting 24 contrasts into a model including the within-subject factors Word Type (4 levels: MO, EMneg, EMpos, NEU) and Word Contrast (6 levels: Empty, PT-2, PT-1, PT, PT+1, Full), with dependence and equal variance assumed between the levels of both factors. Then, we investigated whether the Word Contrast level modulated the [MO – (EMneg+EMpos)] and [(EMneg+EMpos) – MO] activation effects, following either a subliminal (weights: [-5 +1 +1 +1 +1 +1]; order: [Empty PT-2 PT-1 PT PT+1 Full]), a linear [-2.5 -1.5 -0.5 +0.5 +1.5 +2.5] or a stepwise [-1 -1 -1 +1 +1 +1] BOLD amplitude model.

##### 2.6.2.2. Univariate statistical analysis as a function of PAS

To test the MO and EM effects without subjective awareness, but independently of Word Contrast, we further analyzed BOLD responses as a function of PAS ratings. Specifically, for each type of words, all trials in which the reported PAS score was either 1 or 2 (PAS{1,2}) were modelled as indicating no subjective awareness, whereas all trials in which the PAS score was either 3 or 4 (PAS{3,4}) were modelled as indicating subjective awareness. We restricted this analysis to only those subjects that produced comparable response frequencies across all PAS levels. To exclude participants who did not satisfy this criterion, we applied the same procedure as in Tettamanti et al. (2017): first we estimated the maximum likelihood of the individual Contrast-to-PAS linear function to a Contrast-to-PAS linear model function with R. Then we ranked participants based on the resulting Akaike information criterion. None of the participants markedly departed from a linear increase of PAS score with increasing contrast level. However, one participant was excluded from the following analyses because of an exceedingly low number of PAS{3,4} responses, even for the PT+1 and Full contrast levels (see Supplementary Figure 1). Thus, these analyses were performed on a sub-sample of 20 participants.

At the first level, we specified a general linear model with the same parameters as defined above for the analysis as a function of word contrast. However, here we modelled the six runs by assigning the CFS trial events on the basis of PAS score and including the Empty baseline condition. Specifically, each run included 5 regressors of interest: Empty, MOPAS{1,2}, MOPAS{3,4}, EMPAS{1,2}, EMPAS{3,4}, with EM being either EMneg, EMpos, or NEU depending on the run. All the confound regressors were defined as in the analysis as a function of contrast (see above). Within the estimated first-level general linear model, we defined a set of 12 condition-specific contrasts by specifying a weight of +1 for the regressor of interest (e.g., MOPAS{1,2}) and a weight of 0 for all the other regressors.

For the second-level, random-effects full factorial design, we entered the 12 condition- specific contrasts of each participant into a model including the within-subject factors Word Type (4 levels: MO, EMneg, EMpos, NEU) and PAS (3 levels: Empty, PAS{1,2}, PAS{3,4}), with dependence and equal variance assumed between the levels of both factors. In this full factorial design, we investigated whether the absence or presence of subjective perceptual awareness (as reflected by the PAS score) modulated the activation effects of [MO – (EMneg+EMpos)] and [(EMneg+EMpos) – MO], following a subliminal (weights: [-2 +1 +1]; order: [Empty PAS{1,2} PAS{3,4}]) or a stepwise [-1 -1 2] BOLD amplitude model.

#### 2.6.3. Multivariate pattern analyses (MVPA)

MVPA was carried out using the PyMVPA 2.6.5 toolbox (www.pymvpa.org) running under Python 3.9.2. We first re-estimated the SPM12 univariate first-level analyses on preprocessed but spatially unsmoothed fMRI data. We then entered the resulting spmT map images into the MVPA (Misaki et al., 2010). All the included spmT maps incorporated the Empty baseline subtraction (e.g., [MOPT-2 – Empty]). We applied no Z-scoring or averaging within the PyMVPA toolbox. We restricted the analysis to voxels with gray matter tissue probability > .1, which were comprised within 12-mm radius spherical regions of interest in the MOVN and EMLN (Fig. 1A).

We trained a Gaussian Naive Bayes classifier on spmT maps to distinguish the levels of the Word Type factor (MO, EMneg, EMpos) either between all three levels (3 x 3 classification problems) or between pairwise combinations of these levels (2 x 2 classification problems), based on a leave-one-subject-out cross-validation procedure, analogous to a second-level, random effects analysis (Ghio et al., 2016). More specifically, we performed a searchlight analysis (Kriegeskorte et al., 2006) with 4-mm radius spheres, and for each sphere the mean classification accuracy, as well as the confusion matrix of predicted against actual classes, were calculated across leave-one-subject-out folds.

A Monte Carlo procedure was applied to assess the significance of the classification accuracies. In particular, the spmT map labels were randomly permuted over the factor levels in each sphere 10000 times (Stelzer et al., 2013), and the actual classification accuracy was then compared against the random permutation distribution with a declared P < .0001 threshold.

The mean classification accuracies, together with the corresponding confusion matrices are reported for the significant searchlight spheres, resulting from the analysis of 2 different spmT map sets. A first set entered the 3 x 3 and 2 x 2 multivariate pattern analyses as a function of Word Contrast, with each classification problem assessed: i) across the 5 Word Contrast levels (PT-2, PT- 1, PT, PT+1, Full); ii) across levels below perceptual threshold (PT-2, PT-1); iii) across levels above perceptual threshold (PT+1, Full). The second set entered the 3 x 3 and 2 x 2 multivariate pattern analyses as a function of PAS ratings, with each classification problem assessed: i) across all PAS scores (PAS{1,2}, PAS{3,4}); ii) across PAS{1,2}; ii) across PAS{3,4}.

For the 2 x 2 classification problems, we assessed the average spmT value across the voxels of any significant searchlight spheres, and counted in each of the regions of interest within the MOVN and the EMLN the presence of spheres whose average estimate reflected a greater activation for condition A (e.g., MO) minus B (e.g., EMneg), or vice versa for condition B minus A (note that each region of interest may contain spheres with average activation in just one of these two directions, in both, or no significant spheres at all).

## 3. Results

### 3.1. Behavioural results

The Generalized Linear Mixed Model analysis on the PAS behavioural responses during the fMRI session showed that the Word Contrast level significantly modulated the level of subjective awareness, χ2(4) = 2286.3, p = 2.2 x 10-16 (Fig. 2A). Post-hoc multiple comparison tests (Tuckey’s corrected) revealed that every pairwise difference between levels of Word Contrast was significant (all p <.001). The Word Type main effect (χ2(3) = 4.107, p = 0.250) and the Word Type by Word Contrast interaction (χ2(12) = 0.847, p = 1) were not significant predictors of the PAS scores.

**Figure 2.**
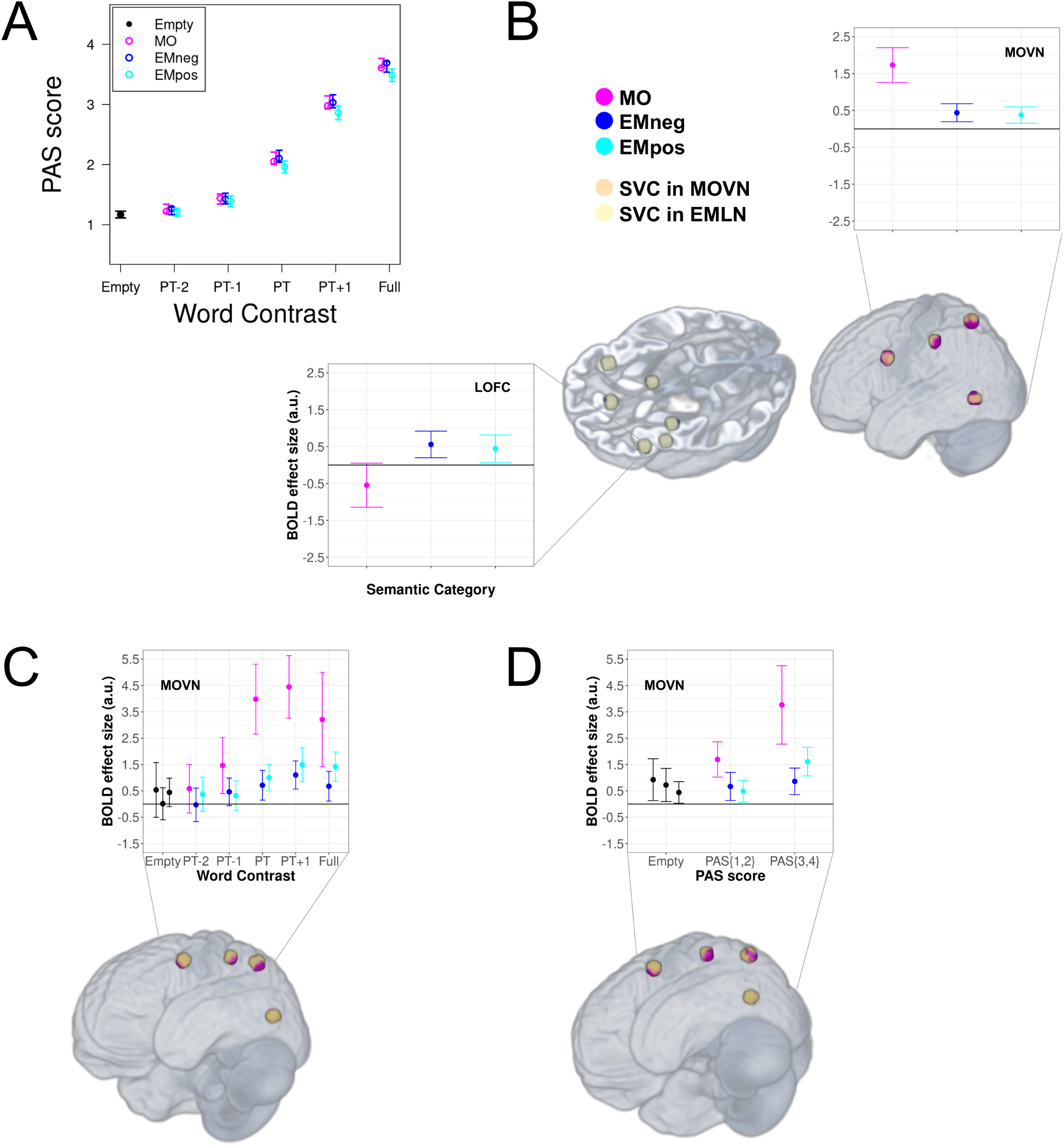
BOLD as a function of Word Type, Word Contrast, or PAS rating. A) Mean (n = 21) behavioral PAS rating scores for the experimental word types as a function of Word Contrast level. Vertical bars indicate 95 % confidence intervals. B) Word Type-specific brain activation in, respectively, the MOVN and the LOFC (as one of the significantly modulated EMLN brain regions). Significant (*p* < .05, small volume FWE correction) brain activations are overlaid (MO: purple clusters; EMneg/EMpos: blue clusters) on the small volumes of interest (MOVN: orange spheres; EMLN: yellow spheres) and displayed on a volumetric rendering of the average anatomical image of all participants. Dot-plots illustrate mean BOLD responses across all significant voxels. Vertical bars indicate 95 % confidence intervals. C) Subliminal MO-specific brain activation in the MOVN as a function of Word Contrast (all conventions are identical to panel B). D) Subliminal MO-specific brain activation in the MOVN as a function of PAS rating (all conventions are identical to panel B).

### 3.2. Univariate fMRI analyses

#### 3.2.1. Effects of Word Type

We found that processing MO elicited greater activations than EM (i.e., EMneg+EMpos) in all the regions of the MOVN (Table 2; Fig. 2B). In turn, when considering EM against MO, we found activations in the sACC, the left OFC, the LAMY and, at trend level, the RAMY (Table 2; Fig. 2B). We found no specific activation for neither the simple nor the quadratic effect of valence, which therefore were not further investigated.

**Table 2.** Effects of Word Type.

|  | Voxels | *p* value | Z score | x, y, z (mm) |
| --- | --- | --- | --- | --- |
| **MO > [EMneg+EMpos]** |  |  |  |  |
| LVPM | 57 | <.001 | 6.11 | -46 8 28 |
| LIPL | 63 | <.001 | 6.23 | -42 -34 44 |
| LSPL | 57 | <.001 | 5.86 | -24 -64 58 |
| LMTG | 46 | .003 | 4.95 | -46 -62 -6 |
| **[EMneg+EMpos] > MO** |  |  |  |  |
| LAMY | 6 | .012 | 3.96 | -22 -8 -16 |
| RAMY | 1 | .042* | 3.60 | 24 0 -14 |
| LOFC | 3 | .010 | 4.02 | -44 26 -10 |
| sACC | 2 | .005 | 4.20 | 2 42 10 |

*Note to Table 2.* MO = manipulable object words; EMneg = emotion words with negative valence; EMpos = emotion words with positive valence; LVPM = left ventral premotor cortex; LIPL= left inferior parietal lobule; LSPL = left superior parietal lobule; LMTG = left middle temporal gyrus; LAMY = left amygdala; RAMY = right amygdala, LOFC = left orbito-frontal cortex; sACC = sub-/pre-genual anterior cingulate cortex; x, y, z cartesian values in mm referring to the MNI space.
\*Trend level, assuming a Bonferroni correction and an alpha = .025.

#### 3.2.2. Effects of Word Type as a function of Word Contrast

We examined whether the levels of Word Contrast modulated activations within the MOVN and the EMLN following a subliminal profile (i.e., activation both above and below perceptual threshold), according to our experimental hypothesis. For comparison, we also tested a linear (i.e., gradual activation increase with contrast) and a stepwise (i.e., activation only above perceptual threshold) BOLD amplitude model.

Subliminal BOLD modulations for MO compared to EM were found in three regions of the MOVN, i.e., LVPM, LIPL, LSPL (Table 3; Fig. 2C). Linear and stepwise modulations for MO compared to EM were found in all regions of the MOVN (Table 3). In turn, we did not find any subliminal, linear, nor stepwise BOLD modulations for EM compared to MO.

**Table 3.** Effects of Word Type as a function of Word Contrast.

|  | Voxels | *p* value | Z score | x, y, z (mm) |
| --- | --- | --- | --- | --- |
| **A. Subliminal** |  |  |  |  |
| **MO > [EMneg+EMpos]** |  |  |  |  |
| LVPM | 15 | .001 | 4.51 | -46 8 28 |
| LIPL | 23 | .005 | 4.12 | -42 -36 44 |
| LSPL | 34 | .002 | 4.31 | -24 -64 58 |
| **B. Linear** |  |  |  |  |
| **MO > [EMneg+EMpos]** |  |  |  |  |
| LVPM | 71 | <.001 | 6.72 | -46 8 28 |
| LIPL | 47 | <.001 | 5.77 | -44 -36 42 |
| LSPL | 57 | <.001 | 4.70 | -22 -62 56 |
| LMTG | 37 | <.001 | 5.88 | -44 -66 -6 |
| **C. Stepwise** |  |  |  |  |
| **MO > [EMneg+EMpos]** |  |  |  |  |
| LVPM | 84 | <.001 | 6.40 | -46 8 28 |
| LIPL | 60 | <.001 | 6.82 | -44 -36 42 |
| LSPL | 71 | <.001 | 5.59 | -28 -60 62 |
| LMTG | 24 | <.001 | 5.70 | -44 -66 -6 |

*Note to Table 3.* MO = manipulable object words; EMneg = emotion words with negative valence; EMpos = emotion words with positive valence; LVPM = left ventral premotor cortex; LIPL= left inferior parietal lobule; LSPL = left superior parietal lobule; LMTG = left middle temporal gyrus; x, y, z cartesian values in mm referring to the MNI space.

#### 3.2.3. Effects of Word Type as a function of PAS

To test the presence of Word Type effects in the absence of subjective awareness, independently of Word Contrast, we further analyzed BOLD responses as a function of PAS ratings. We tested whether activations within the regions of interest conformed to a subliminal and/or a stepwise BOLD amplitude model.

We found stronger subliminal and stepwise BOLD modulations for MO compared to EM in all regions of the MOVN (Table 4; Fig. 2D; Supplementary Figure 1). We did not found any stronger subliminal nor stepwise BOLD modulations for EM compared to MO.

**Table 4.** Effects of Word Type as a function of PAS score.

|  | Voxels | *p* value | Z score | x, y, z (mm) |
| --- | --- | --- | --- | --- |
| **A. Subliminal** |  |  |  |  |
| **MO > [EMneg+EMpos]** |  |  |  |  |
| LVPm | 39 | <.001 | 5.58 | -46 8 28 |
| LIPL | 46 | .001 | 4.51 | -44 -34 44 |
| LSPL | 24 | .002 | 4.02 | -24 -64 60 |
| LMTG | 1 | .001 | 3.62 | -46 -68 -6 |
| **B. Stepwise** |  |  |  |  |
| **MO > [EMneg+EMpos]** |  |  |  |  |
| LVPm | 82 | <.001 | 8.0 | -46 8 28 |
| LIPL | 81 | <.001 | 5.85 | -44 -36 44 |
| LSPL | 34 | .001 | 4.52 | -28 -60 62 |
| LMTG | 33 | <.001 | 5.13 | -46 -68 -6 |

*Note to Table 4.* MO = manipulable object words; EMneg = emotion words with negative valence; EMpos = emotion words with positive valence; LVPm = left ventral premotor cortex; LIPL= left inferior parietal lobule; LSPL = left superior parietal lobule; LMTG = left middle temporal gyrus; LAMY = left amygdala; x, y, z cartesian values in mm referring to the MNI space.

### 3.3. Multivariate classification of fMRI activation patterns

To evaluate whether activation differences between MO and EM, as resulting from the univariate fMRI analyses, reflected distinct category-specific neural codes that could be distinguished by a machine learning classifier, we also ran a searchlight MVPA on the same fMRI data.

#### 3.3.1. MVPA as a function of Word Contrast

In the 3 x 3 analyses as a function of Word Contrast (Fig. 3A), reflecting the main effect of Word Type, significant spheres distinguishing among the 3 types of words with accuracy above chance (chance level = 33.3%) were found, either when we considered all Word Contrast levels (173 spheres distributed over all regions of interest except LAMY; mean classification accuracy = 44.4%), just those above PT (36 spheres distributed over RAMY, LOFC, ROFC, LINS, sACC, LVPM, LIPL, LSPL, LMTG; mean accuracy = 49.8%), or, most importantly, just the Word Contrast levels below PT (3 spheres over ROFC and LINS; mean accuracy = 50.8%).

**Figure 3.**
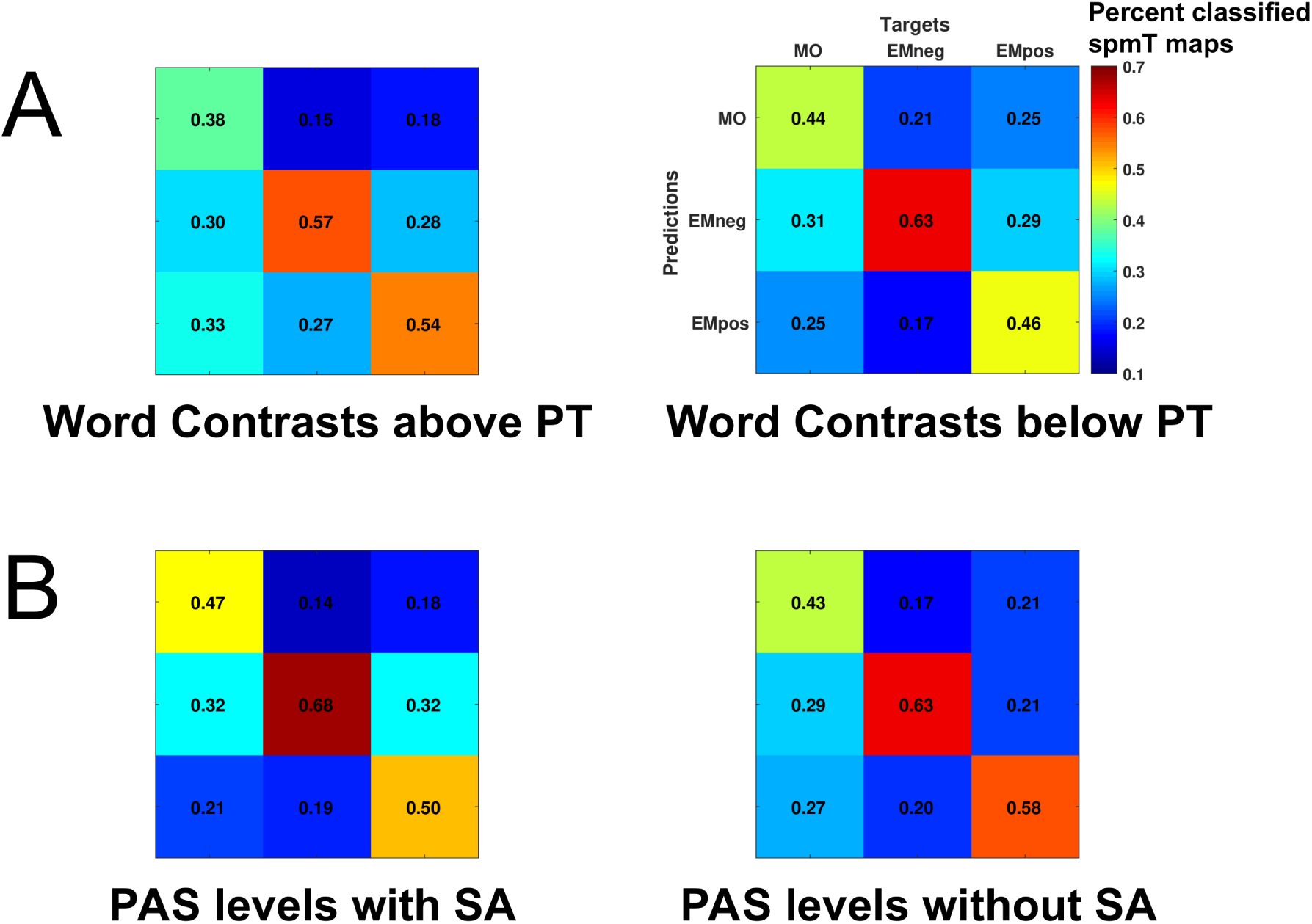
Multivariate pattern analysis. Confusion matrices (labels for all matrices correspond to those displayed on the top right matrix) representing the percent classification accuracy across all significant (*p* < .0001; 10000 permutations) searchlight spheres. A) 3 x 3 classification of Word Type as a function of Word Contrast for all levels above (left) or below perceptual threshold (right). B) 3 x 3 classification of Word Type as a function of PAS rating for all levels indicating presence (left) or absence of subjective awareness (right).

In the 2 x 2 analyses as a function of Word Contrast (Table 5), reflecting the pairwise post- hoc comparisons between the different word types, the MVPA classifier was able to distinguish significantly above chance (chance level = 50%) among all word types pairs, again either when we considered all Word Contrast levels, or just those above or just those below perceptual threshold. Significant classification spheres were variably distributed across the 10 regions of interest, but with a relative greater proportion in regions of the MOVN of spheres with greater activation for MO (particularly the LSPL and LMTG), and a relatively greater proportion of spheres whose average BOLD estimate reflected a greater activation for EM words (EMneg and EMpos) in the EMLN regions of interest.

**Table 5.** MVPA: 2 x 2 analyses as a function of Word Contrast.

| classification | PT | Mean Acc | N | MOVN |  |  |  | EMLN |  |  |  |  |  |
| --- | --- | --- | --- | --- | --- | --- | --- | --- | --- | --- | --- | --- | --- |
|  |  |  |  | LVPM | LSPL | LIPL | LMTG | LAMY | RAMY | LOFC | ROFC | LINS | sACC |
| MO vs EM<sub>neg</sub> | All | 63.4% | 169 | X | X |  | X | X | XX | XX | XX | XX | X |
| MO vs EM<sub>neg</sub> | aPT | 69.4% | 35 | X | XX | X | XX |  | X | X | XX | X | X |
| MO vs EM<sub>neg</sub> | bPT | 71.6% | 11 |  |  |  |  |  |  | X | X | X |  |
| MO vs EM<sub>pos</sub> | All | 62.6% | 110 | X | XX | X | X | X | X | XX | XX | XX |  |
| MO vs EM<sub>pos</sub> | aPT | 68.5% | 17 | XX | X |  | X |  | X |  |  |  |  |
| MO vs EM<sub>pos</sub> | bPT | 70.0% | 5 | X | X |  | X |  |  |  |  |  |  |
| EM<sub>neg</sub> vs EM<sub>pos</sub> | All | 64.0% | 117 | X |  | X | X | X | XX | X | X | XX | X |
| EM<sub>neg</sub> vs EM<sub>pos</sub> | aPT | 70.1% | 9 |  |  | X |  |  | X | X |  | X | X |
| EM<sub>neg</sub> vs EM<sub>pos</sub> | bPT | 71.4% | 1 |  |  |  |  |  |  |  |  | X |  |

*Note to Table 5.* An “X” indicates the successful classification of the pair of word types within the region of interest, with the colour indicating the direction of associated mean spmT values. PT = perceptual threshold, aPT = above perceptual threshold, bPT = below perceptual threshold, N = number of voxels in the cluster. MO = manipulable object words; EM<sub>neg</sub> = emotion words with negative valence; EM<sub>pos</sub> = emotion words with positive valence. MOVN = MO-specific visuomotor network, comprising: LVPM = left ventral premotor cortex, LIPL = left inferior parietal lobule, LSPL = left superior parietal lobule, LMTG = left middle temporal gyrus. EMLN = EM-specific limbic network, comprising: LAMY = left amygdala, RAMY = right amygdala, LOFC = left orbito-frontal cortex, ROFC = right orbito-frontal cortex, LINS = left anterior insula, sACC = sub-/pre-genual anterior cingulate cortex.

#### 3.3.2. MVPA as a function of PAS

Also in the 3 x 3 analyses as a function of PAS ratings (Fig. 3B), the three word types where significantly distinguished above chance (chance level = 33.3%), either when we considered all PAS levels (33 spheres distributed over LOFC, ROFC, LINS, sACC LVPM, LSPL, LMTG; mean classification accuracy = 49.6%), or just those indicating presence of subjective awareness (11 spheres distributed over LOFC, LINS, LVPM, and LSPL; mean accuracy = 55.2%). When we considered just the PAS levels indicating absence of subjective awareness, significant spheres were only found at a lower significance threshold (P < 0.001 against 10000 permutations; 9 spheres over LOFC, ROFC, LINS, sACC, and LMTG; mean accuracy = 54.8%).

In the 2 x 2 analyses as a function of PAS ratings (Table 6), the MVPA classifier distinguished above chance all word types pairs and for all three PAS rating level combinations (all levels, just those indicating presence, or just those indicating absence of subjective awareness), except for the pair EMneg versus EMpos (with and without subjective awareness), which was only distinguished at a lower significance threshold (P < 0.001 against 10000 permutations). Classification spheres were variably distributed across the 10 regions of interest, but with a slightly greater proportion in MOVN regions of spheres with greater activation for MO, and a slightly greater proportion of spheres whose average BOLD estimate reflected a greater activation for EM (EMneg and EMpos) in EMLN regions of interest.

**Table 6.** MVPA: 2 x 2 analyses as a function of PAS.

| classification | SA | Mean Acc | N | MOVN |  |  |  | EMLN |  |  |  |  |  |
| --- | --- | --- | --- | --- | --- | --- | --- | --- | --- | --- | --- | --- | --- |
|  |  |  |  | LVPM | LSPL | LIPL | LMTG | LAMY | RAMY | LOFC | ROFC | LINS | sACC |
| MO vs EM<sub>neg</sub> | All | 70.5% | 20 |  |  | X | X | X X | X |  | X X |  |  |
| MO vs EM<sub>neg</sub> | wSA | 79.6% | 6 | X |  |  |  |  | X | X |  | X |  |
| MO vs EM<sub>neg</sub> | w/oSA | 81.3% | 2 |  |  |  |  |  |  | X |  | X |  |
| MO vs EM<sub>pos</sub> | All | 71.1% | 8 |  |  |  |  |  | X | X | X |  |  |
| MO vs EM<sub>pos</sub> | wSA | 76.0% | 5 |  |  | X | X |  |  | X |  |  |  |
| MO vs EM<sub>pos</sub> | w/oSA | 76.3% | 2 | X X |  |  |  |  |  |  |  |  |  |
| EM<sub>neg</sub> vs EM<sub>pos</sub> | All | 71.0% | 18 |  |  |  | X | X | X | X | X X | X | X |
| EM<sub>neg</sub> vs EM<sub>pos</sub> | wSA | 75.0% | 6 | (X) |  | (X) | (X) |  |  | (X) |  | (X) |  |
| EM<sub>neg</sub> vs EM<sub>pos</sub> | w/oSA | 76.9% | 4 |  |  |  |  |  |  |  | (X)(X) | (X) |  |

*Note to Table 6.* An “X” indicates the successful classification of the pair of word types within the ROI, with the colour indicating the direction of associated mean spmT values. SA = subjective awareness, wSA = with subjective awareness, w/oSA = without subjective awareness, N = number of voxels in the cluster. MO = manipulable object words; EM<sub>neg</sub> = emotion words with negative valence; EM<sub>pos</sub> = emotion words with positive valence. MOVN = MO-specific visuomotor network, comprising: LVPM = left ventral premotor cortex, LIPL= left inferior parietal lobule, LSPL = left superior parietal lobule, LMTG = left middle temporal gyrus. EMLN = EM-specific limbic network, comprising: LAMY = left amygdala, RAMY = right amygdala, LOFC = left orbito-frontal cortex, ROFC = right orbito-frontal cortex, LINS = left anterior insula, sACC = sub-/pre-genual anterior cingulate cortex.

## 4. Discussion

The present fMRI study aimed at testing whether brain regions coding for experience- derived information that constitutes conceptual representations are automatically recruited during the subliminal perception of MO- and EM-related words. Univariate analyses of BOLD signal showed a MO word-specific activation of the MOVN, a left-lateralized visuomotor network involved in object-directed action and object identification (Caspers et al., 2010; Ishibashi et al., 2016), and supposed to represent and process visual and motor-related features such as manipulability and affordances (Buxbaum and Kalénine, 2010; Sakreida et al., 2016). Notably, this activation was found even under subliminal unaware processing conditions, both when the words were presented below perceptual threshold and when participants subjectively reported a lack of stimulus awareness. EM word processing specifically engaged the EMLN, a bilateral limbic network involved in emotion perception and recognition (Dalgleish, 2004; Dahlén et al., 2022), and supposed to represent affective properties (Kober et al., 2008; Lindquist et al, 2012). Univariate analyses, however, did not provide evidence for EM word-specific effects under subliminal unaware processing conditions. MVPA, instead, showed that distinct MO and EM word-specific neural codes could be distinguished above chance in MOVN and EMLN, when words were presented both above and below the perceptual threshold, as well as with and without subjective perceptual awareness.

Previous CFS studies have provided conflicting evidence for the recruitment of the MOVN as a function of awareness when processing tool pictures subliminally: whereas some works found activations in some but not all regions of this left-hemispheric premotor-parietal-temporal network (Fang & He, 2005; Almeida et al., 2010; Suzuki et al., 2014), other studies did not report any significant effects (Hesselmann & Malach, 2011; Hesselmann et al., 2011; Fogelson et al., 2014; Ludwig et al., 2015, 2016). In contrast, we previously showed the recruitment and effective connectivity of the MOVN both under aware and unaware processing of MO pictures (Tettamanti et al., 2017; Ghio et al., 2022). Here, we confirmed and expanded these findings, in line with the hypothesis that semantic features are automatically retrieved also during MO words processing, thus despite the lack of a visual or other sensory depiction of tools, and so excluding the potentially confounding effect of perceptual properties, such as the prototypical elongated shape of MO. Indeed, we found that MO elicited a subliminal BOLD response profile as a function of Word Contrast in LVPM, LIPL, and SPL of the MOVN, that is a stable activation regardless of stimulus perception being under aware or unaware conditions. These three regions also exhibited a linear and a stepwise response profile, in concomitance with the subliminal one. Taken together, the significance of all three modulations indicates activation both above and below perceptual threshold, with a relatively stronger activation with than without awareness, and with relatively higher response amplitudes as the word contrast level increased. A fourth MOVN area, namely LMTG, mainly implied in object identification (Bracci et al., 2012), showed only linear and stepwise haemodynamic modulations. However, this effect does not contradict our hypothesis, as it still suggests a significant activation of this region below the perceptual threshold, although weaker than above. Furthermore, assessing fMRI activation as a function of PAS ratings revealed significant subliminal and stepwise modulations for all MOVN regions, indicting that these regions are activated both in the presence and, although to a lesser extent, in the absence of subjective awareness. Consistently with the univariate analyses, MVPA showed that a supporting vector machine classifier trained on the activity in MOVN regions was able to successfully tell apart MO and EM activation maps, not only above but also below perceptual threshold, and both with and without subjective perceptual awareness. Taken together, our results suggest that subliminal perception of visually presented MO words automatically activate conceptual-semantic decoding of visuomotor and functional MO-related information that is likely encoded during previous experiences with tool exemplars (Binder & Desai, 2011; Lambon Ralph et al., 2017).

Regarding EM word processing, we found that it specifically involved the EMLN, a network engaged in what Lindquist and colleagues (2012) defined as “core affect” domain. This term refers to the mental representations sustaining the semantic processing of discrete emotions, and comprising bodily sensations – somatovisceral, kinesthetic, proprioceptive, and neurochemical – experienced along two main dimensions: valence and arousal. Previous CFS studies have suggested that valence plays an important role in an EM stimulus emergence to consciousness (Jiang & He, 2006; Vizueta et al., 2012), even though evidence is inconsistent. While some studies reported a negativity bias (i.e., negatively-valenced words emerging from suppression faster than positively-valenced and neutral stimuli; Yang & Yeh, 2011; Sklar et al., 2012), other works reported the opposite pattern (Vinson et al., 2011) or no effect of valence (Rabagliati et al., 2018; Cheng et al., 2019). Our univariate analysis revealed no subliminal response profile, either as a function of Word Contrast or as a function of PAS scores, and no parametric effects of valence, either simple or quadratic. However, the entire EMLN was consistently implicated in the multivariate searchlight decoding of EMneg versus EMpos activation patterns, both when considering contrast levels below perceptual threshold and the absence of subjective awareness. Considering the theoretical and methodological differences between univariate and multivariate analysis (Hebart & Baker, 2018) which make the latter more sensitive to detecting mental representations (Norman et al., 2006), our findings suggest that voxels within EMLN regions are likely not more activated for positively than negatively valenced EM words (or vice versa), but rather exhibit patterns of distributed activations or de-activations which give rise to a clearly identifiable EM-specific neural coding in the EMLN, even under unaware processing.

Our results about the subliminal activation of the MOVN and the EMLN suggest that unaware processing of visually presented MO and EM words automatically elicits the retrieval of, respectively, tool-related visuomotor properties (e.g., features useful to identify objects as well as to functionally interact with them, such as manipulability and affordance) and core affect (e.g., interoceptive and somatic) information. Since (i) no overt task implying, for instance, imagery or deep semantic processing, was required to participants, and (ii) neither operationalized nor subjective awareness was implicated in word processing, we suggest that the category-specific features encoded by MOVN and EMLN regions are constitutively integrated into the conceptual representation of MO and EM. These findings are consistent with grounded cognition theories of conceptual knowledge, according to which concepts rely, at least partially, on the reuse of specific information derived by previous experience with similar percepts (Meteyard et al., 2012; Lambon Ralph et al., 2017; Kiefer & Harpaintner, 2020). This information refers to sensory-motor experiences in the case of MO (see Willems & Casasanto, 2011), and to internal (i.e., proprioceptive, somatovisceral) states in the case of EM (Winkielman et al., 2018). The hypothesis that these category-specific features are encoded over multiple experiences of EM and MO instances, represented in conceptual representations, and recruited every time the corresponding concepts are semantically processed, is consistent with the view of the brain as a predictive system, evolved to efficiently meet the ongoing needs of the body (Adams et al., 2013; Clark, 2013). Whether the circumstances require to promptly interact with physical objects outside the body, or to cope with physiological states inside the body, knowing how to plan an appropriate response is essential for survival (Frye & Lindquist, in press). In this scenario, language plays an important role: words rapidly and automatically activate properties of the entities they refer to, which overlap with those elicited during the perception of and the experience with such entities. By considering a broader perspective on language and semantic processing, this mechanism can be considered in line with the idea suggested by Lupyan and colleagues that language acts as a flexible “artificial” context, according to which the brain unconsciously generates predictive beliefs about incoming inputs, possibly through a fast re-tuning of the organism’s entire semantic network , thereby helping constraining which type of top-down information is useful to recruit and what impact it has on lower-level processes as well as on inference and reasoning (Lupyan & Ward, 2013; Lupyan & Clark, 2015).

In conclusion, previous evidence has demonstrated that visually perceiving objects in the absence of awareness elicits a comparable neurophysiological response signature to that evoked by their conscious perception, thus reflecting not only a visual but also a semantic processing (Fahrenfort et al., 2017; Tettamanti et al., 2017). We extend these findings by providing evidence that also tool words, which are arbitrarily associated with their referents through learning and do not carry any object-related visuomotor information, recruit a manipulable object-specific visuomotor network. Additionally, we generalize these results to the domain of discrete emotions, by demonstrating that processing emotion label words activates portions of the core affect limbic network. Crucially, MO and EM specific neural activations are decoded even in the absence of perceptual awareness, suggesting that experience-derived information constituting conceptual representations is automatically recruited.

## Data and Code Availability

The data and analysis code are available at https://doi.org/10.17632/gfb3xb5yty.1.

## Author contributions

M.G., B.C., and M.T. designed the experiment; M.G. and B.C. collected the data; M.G, B.C., M.T. analyzed the data; M.G., B.C., M.T. organized and wrote the manuscript.

---

**Supplementary Figure 1.**
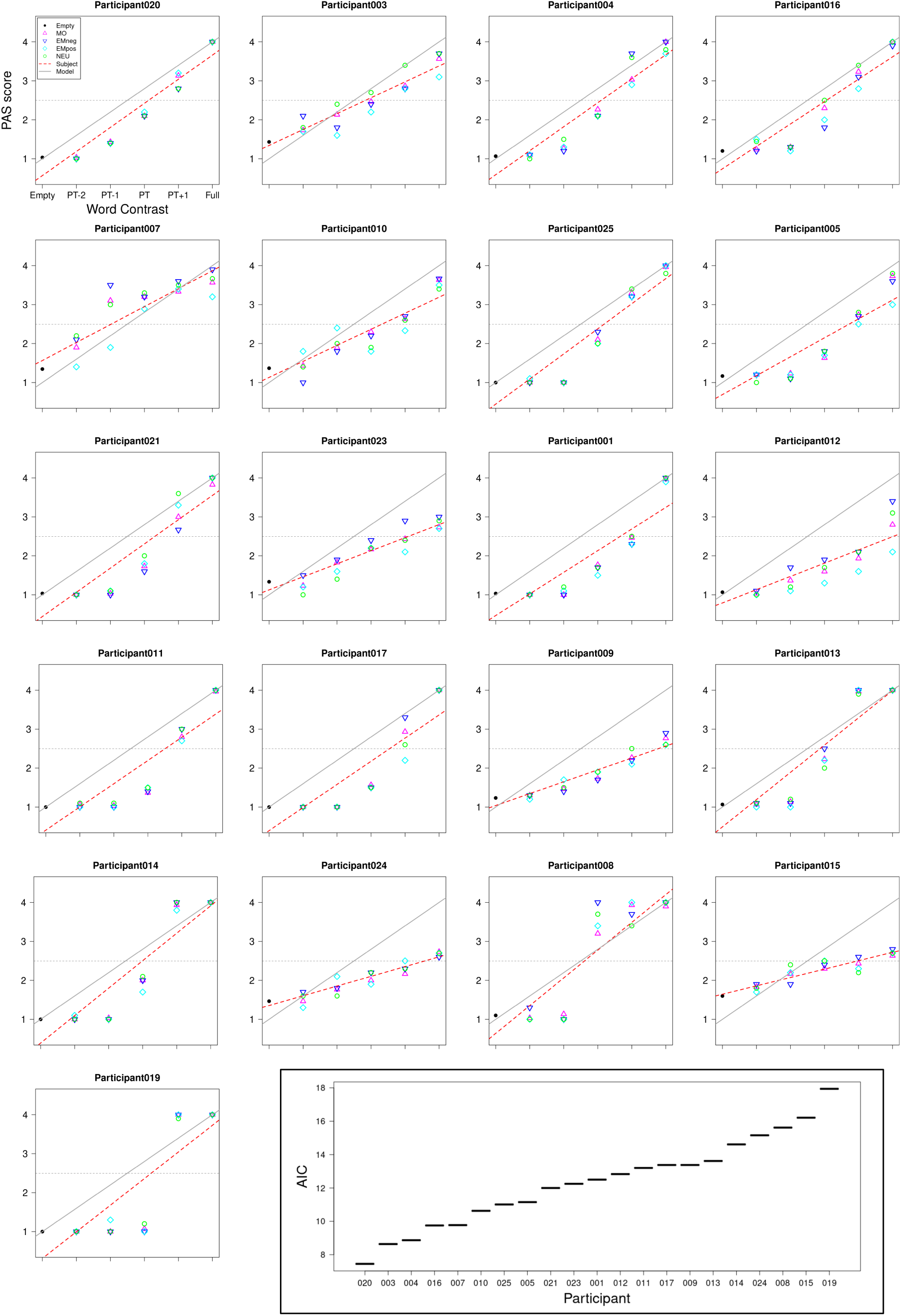
Selection of participants for the BOLD analysis as a function of PAS. The participants were ranked according to the Akaike information criterion (AIC; bottom right inset), obtained by Maximum Likelihood Estimation of the fit of the individual Word Contrast-to-PAS linear function (red dashed line; calculated on the average PAS ratings across all stimulus types (Empty, MO, EMneg, EMpos, NEU)) to a Word Contrast-to-PAS linear model function (gray line). The conventions of the dot-plots of all participants are as displayed for the “Participant020” on top left. None of the participants markedly departed from a linear increase of PAS score with increasing contrast level. However, “Participant012” was excluded from the analyses of BOLD activation as a function of PAS rating, because of an exceedingly low number of PAS{3,4} responses. This left the sub-sample for these analyses with 20 participants.

**Supplementary Table 1A.**
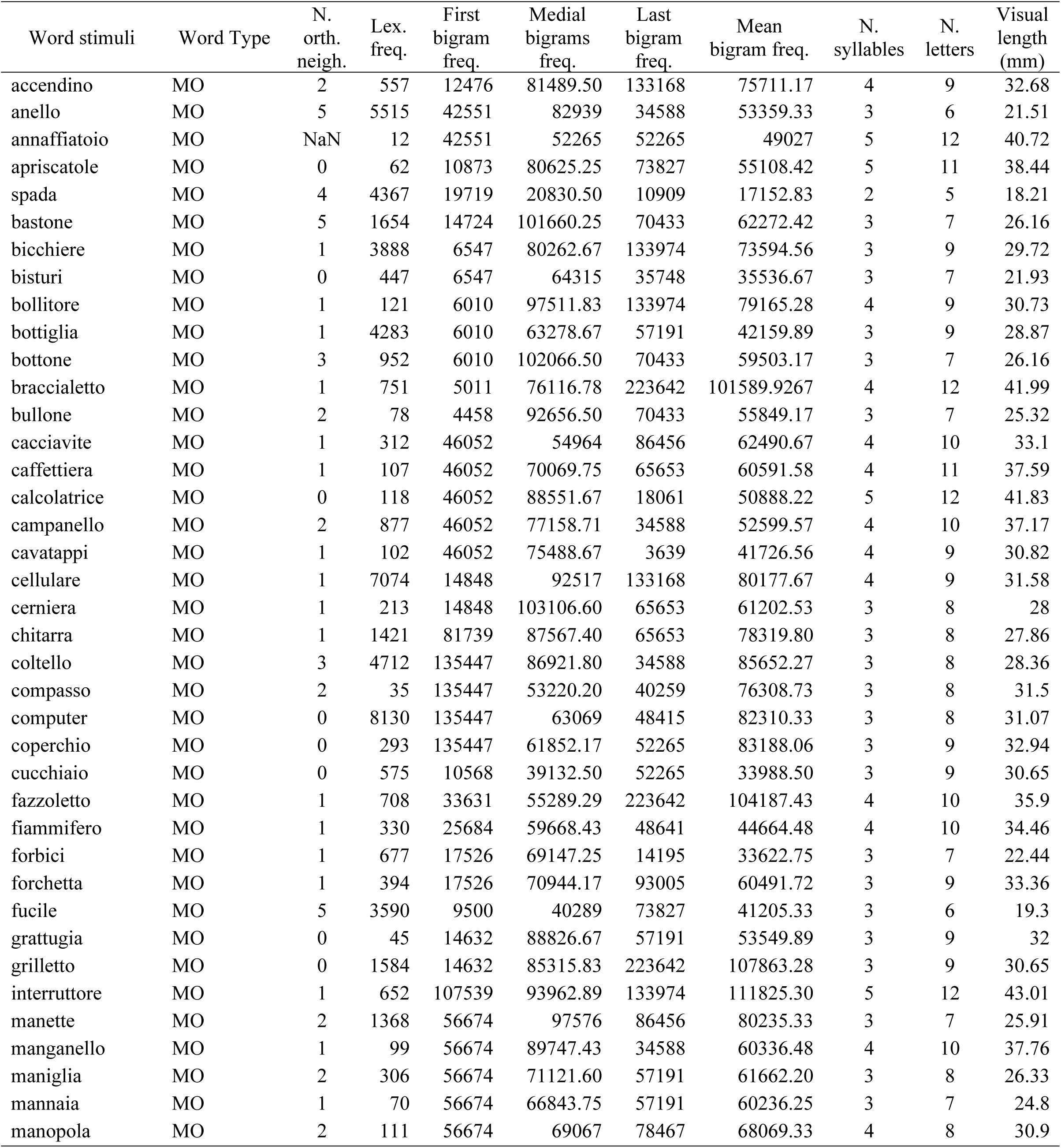
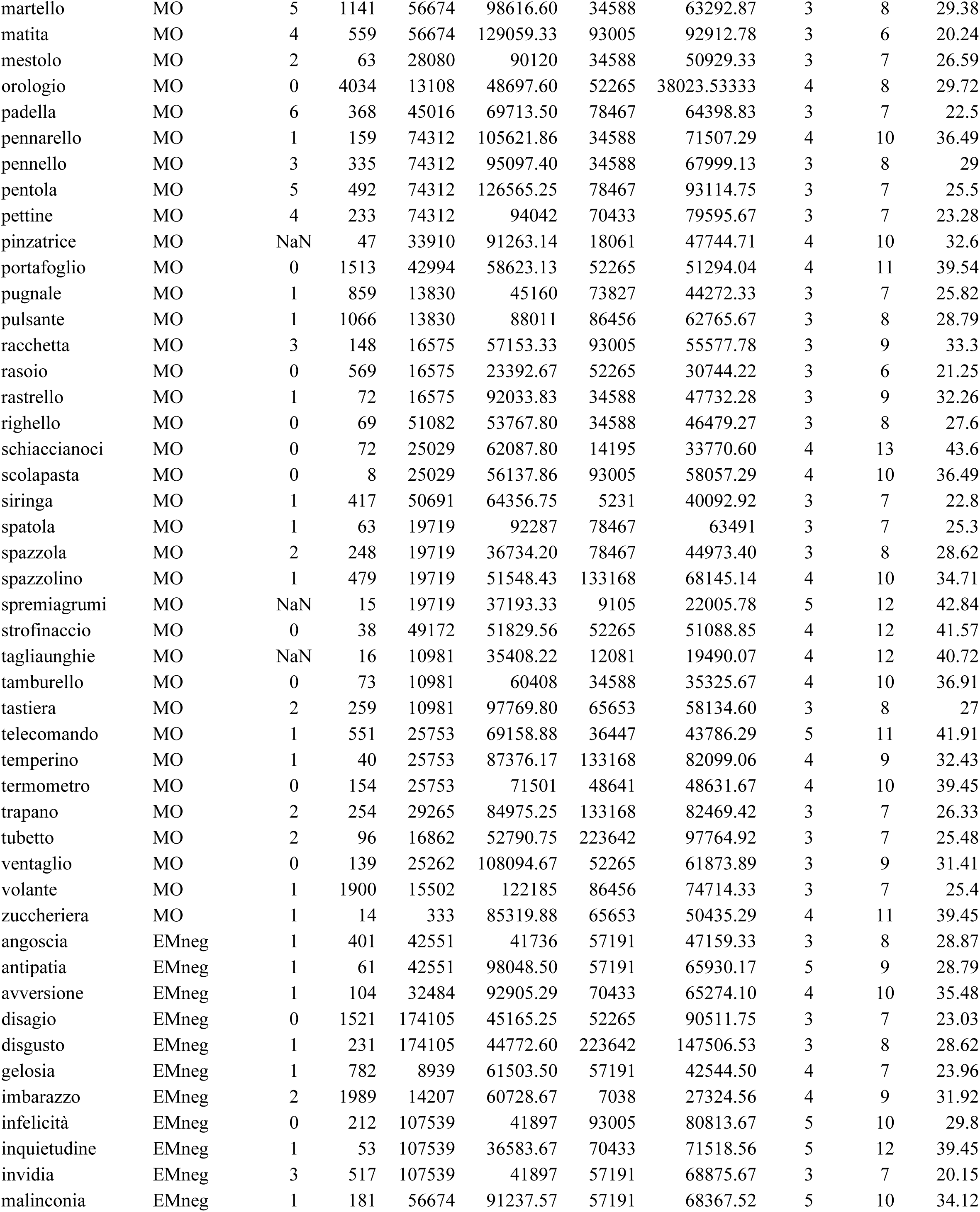
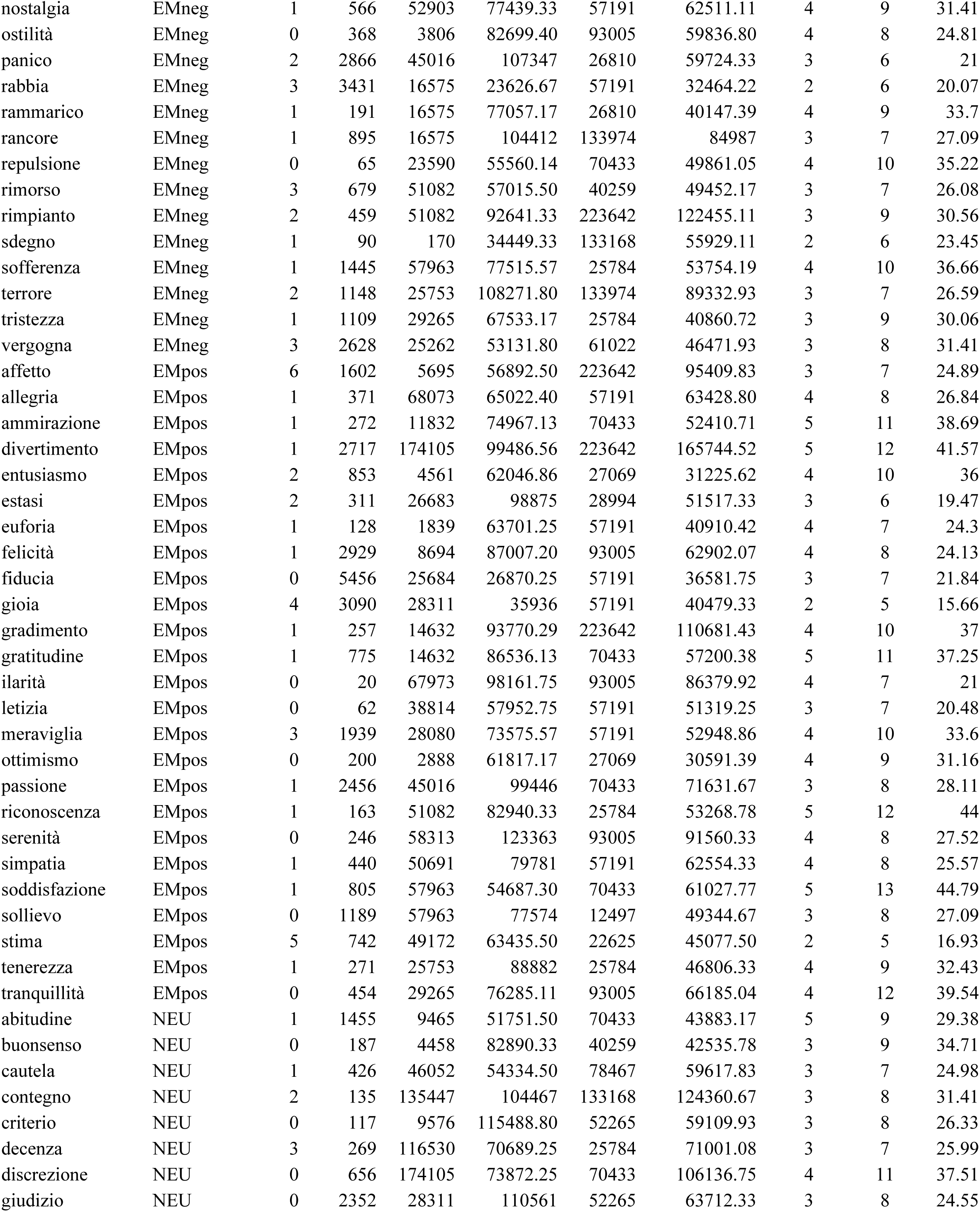
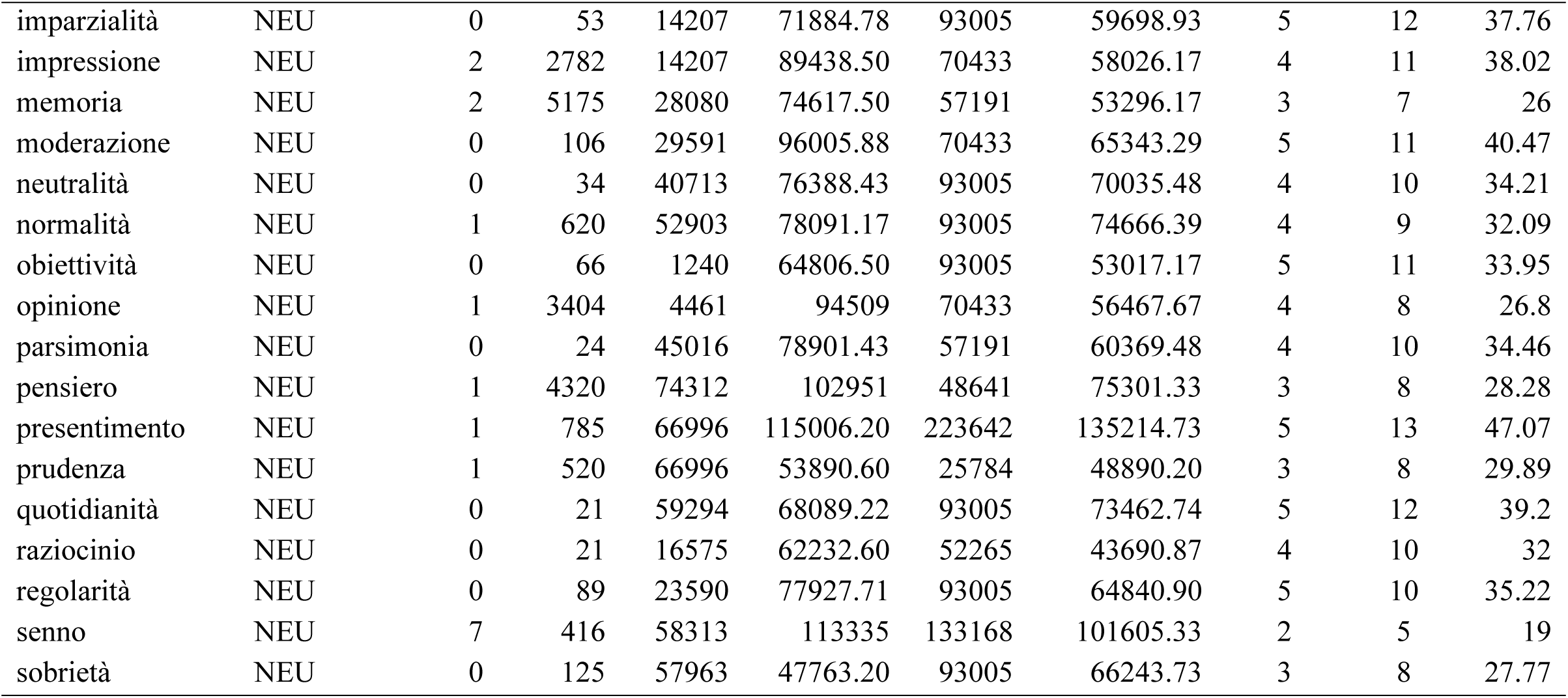
Lexical and sublexical variables of experimental stimuli.

| Word stimuli | Word Type | N.<br>orth.<br>neigh. | Lex.<br>freq. | First<br>bigram<br>freq. | Medial<br>bigrams<br>freq. | Last<br>bigram<br>freq. | Mean<br>bigram freq. | N.<br>syllables | N.<br>letters | Visual<br>length<br>(mm) |
| --- | --- | --- | --- | --- | --- | --- | --- | --- | --- | --- |
| accendino | MO | 2 | 557 | 12476 | 81489.50 | 133168 | 75711.17 | 4 | 9 | 32.68 |
| anello | MO | 5 | 5515 | 42551 | 82939 | 34588 | 53359.33 | 3 | 6 | 21.51 |
| annaffiatoio | MO | NaN | 12 | 42551 | 52265 | 52265 | 49027 | 5 | 12 | 40.72 |
| apricatole | MO | 0 | 62 | 10873 | 80625.25 | 73827 | 55108.42 | 5 | 11 | 38.44 |
| spada | MO | 4 | 4367 | 19719 | 20830.50 | 10909 | 17152.83 | 2 | 5 | 18.21 |
| bastone | MO | 5 | 1654 | 14724 | 101660.25 | 70433 | 62272.42 | 3 | 7 | 26.16 |
| bicchiera | MO | 1 | 3888 | 6547 | 80262.67 | 133974 | 73594.56 | 3 | 9 | 29.72 |
| bisturi | MO | 0 | 447 | 6547 | 64315 | 35748 | 35536.67 | 3 | 7 | 21.93 |
| bollitore | MO | 1 | 121 | 6010 | 97511.83 | 133974 | 79165.28 | 4 | 9 | 30.73 |
| bottiglia | MO | 1 | 4283 | 6010 | 63278.67 | 57191 | 42159.89 | 3 | 9 | 28.87 |
| bottone | MO | 3 | 952 | 6010 | 102066.50 | 70433 | 59503.17 | 3 | 7 | 26.16 |
| braccialetto | MO | 1 | 751 | 5011 | 76116.78 | 223642 | 101589.9267 | 4 | 12 | 41.99 |
| bullone | MO | 2 | 78 | 4458 | 92656.50 | 70433 | 55849.17 | 3 | 7 | 25.32 |
| cacciavite | MO | 1 | 312 | 46052 | 54964 | 86456 | 62490.67 | 4 | 10 | 33.1 |
| caffettiera | MO | 1 | 107 | 46052 | 70069.75 | 65653 | 60591.58 | 4 | 11 | 37.59 |
| calcolatrice | MO | 0 | 118 | 46052 | 88551.67 | 18061 | 50888.22 | 5 | 12 | 41.83 |
| campanello | MO | 2 | 877 | 46052 | 77158.71 | 34588 | 52599.57 | 4 | 10 | 37.17 |
| cavatappi | MO | 1 | 102 | 46052 | 75488.67 | 3639 | 41726.56 | 4 | 9 | 30.82 |
| cellulare | MO | 1 | 7074 | 14848 | 92517 | 133168 | 80177.67 | 4 | 9 | 31.58 |
| cerniera | MO | 1 | 213 | 14848 | 103106.60 | 65653 | 61202.53 | 3 | 8 | 28 |
| chitarra | MO | 1 | 1421 | 81739 | 87567.40 | 65653 | 78319.80 | 3 | 8 | 27.86 |
| coltello | MO | 3 | 4712 | 135447 | 86921.80 | 34588 | 85652.27 | 3 | 8 | 28.36 |
| compasso | MO | 2 | 35 | 135447 | 53220.20 | 40259 | 76308.73 | 3 | 8 | 31.5 |
| computer | MO | 0 | 8130 | 135447 | 63069 | 48415 | 82310.33 | 3 | 8 | 31.07 |
| coperchio | MO | 0 | 293 | 135447 | 61852.17 | 52265 | 83188.06 | 3 | 9 | 32.94 |
| cucchiaio | MO | 0 | 575 | 10568 | 39132.50 | 52265 | 33988.50 | 3 | 9 | 30.65 |
| fazzoletto | MO | 1 | 708 | 33631 | 55289.29 | 223642 | 104187.43 | 4 | 10 | 35.9 |
| fiammifero | MO | 1 | 330 | 25684 | 59668.43 | 48641 | 44664.48 | 4 | 10 | 34.46 |
| forbici | MO | 1 | 677 | 17526 | 69147.25 | 14195 | 33622.75 | 3 | 7 | 22.44 |
| forchetta | MO | 1 | 394 | 17526 | 70944.17 | 93005 | 60491.72 | 3 | 9 | 33.36 |
| fucile | MO | 5 | 3590 | 9500 | 40289 | 73827 | 41205.33 | 3 | 6 | 19.3 |
| grattugia | MO | 0 | 45 | 14632 | 88826.67 | 57191 | 53549.89 | 3 | 9 | 32 |
| grilletto | MO | 0 | 1584 | 14632 | 85315.83 | 223642 | 107863.28 | 3 | 9 | 30.65 |
| interruttore | MO | 1 | 652 | 107539 | 93962.89 | 133974 | 111825.30 | 5 | 12 | 43.01 |
| manette | MO | 2 | 1368 | 56674 | 97576 | 86456 | 80235.33 | 3 | 7 | 25.91 |
| manganello | MO | 1 | 99 | 56674 | 89747.43 | 34588 | 60336.48 | 4 | 10 | 37.76 |
| maniglia | MO | 2 | 306 | 56674 | 71121.60 | 57191 | 61662.20 | 3 | 8 | 26.33 |
| mannaia | MO | 1 | 70 | 56674 | 66843.75 | 57191 | 60236.25 | 3 | 7 | 24.8 |
| manopola | MO | 2 | 111 | 56674 | 69067 | 78467 | 68069.33 | 4 | 8 | 30.9 |
| martello | MO | 5 | 1141 | 56674 | 98616.60 | 34588 | 63292.87 | 3 | 8 | 29.38 |
| matita | MO | 4 | 559 | 56674 | 129059.33 | 93005 | 92912.78 | 3 | 6 | 20.24 |
| mestolo | MO | 2 | 63 | 28080 | 90120 | 34588 | 50929.33 | 3 | 7 | 26.59 |
| orologio | MO | 0 | 4034 | 13108 | 48697.60 | 52265 | 38023.53333 | 4 | 8 | 29.72 |
| padella | MO | 6 | 368 | 45016 | 69713.50 | 78467 | 64398.83 | 3 | 7 | 22.5 |
| pennarello | MO | 1 | 159 | 74312 | 105621.86 | 34588 | 71507.29 | 4 | 10 | 36.49 |
| pennello | MO | 3 | 335 | 74312 | 95097.40 | 34588 | 67999.13 | 3 | 8 | 29 |
| pentola | MO | 5 | 492 | 74312 | 126565.25 | 78467 | 93114.75 | 3 | 7 | 25.5 |
| pettine | MO | 4 | 233 | 74312 | 94042 | 70433 | 79595.67 | 3 | 7 | 23.28 |
| pinzatrice | MO | NaN | 47 | 33910 | 91263.14 | 18061 | 47744.71 | 4 | 10 | 32.6 |
| portafoglio | MO | 0 | 1513 | 42994 | 58623.13 | 52265 | 51294.04 | 4 | 11 | 39.54 |
| pugnale | MO | 1 | 859 | 13830 | 45160 | 73827 | 44272.33 | 3 | 7 | 25.82 |
| pulsante | MO | 1 | 1066 | 13830 | 88011 | 86456 | 62765.67 | 3 | 8 | 28.79 |
| racchetta | MO | 3 | 148 | 16575 | 57153.33 | 93005 | 55577.78 | 3 | 9 | 33.3 |
| rasoio | MO | 0 | 569 | 16575 | 23392.67 | 52265 | 30744.22 | 3 | 6 | 21.25 |
| rastrello | MO | 1 | 72 | 16575 | 92033.83 | 34588 | 47732.28 | 3 | 9 | 32.26 |
| righello | MO | 0 | 69 | 51082 | 53767.80 | 34588 | 46479.27 | 3 | 8 | 27.6 |
| schiaccianoci | MO | 0 | 72 | 25029 | 62087.80 | 14195 | 33770.60 | 4 | 13 | 43.6 |
| scolapasta | MO | 0 | 8 | 25029 | 56137.86 | 93005 | 58057.29 | 4 | 10 | 36.49 |
| siringa | MO | 1 | 417 | 50691 | 64356.75 | 5231 | 40092.92 | 3 | 7 | 22.8 |
| spatola | MO | 1 | 63 | 19719 | 92287 | 78467 | 63491 | 3 | 7 | 25.3 |
| spazzola | MO | 2 | 248 | 19719 | 36734.20 | 78467 | 44973.40 | 3 | 8 | 28.62 |
| spazzolino | MO | 1 | 479 | 19719 | 51548.43 | 133168 | 68145.14 | 4 | 10 | 34.71 |
| spremiagrumi | MO | NaN | 15 | 19719 | 37193.33 | 9105 | 22005.78 | 5 | 12 | 42.84 |
| strofinaccio | MO | 0 | 38 | 49172 | 51829.56 | 52265 | 51088.85 | 4 | 12 | 41.57 |
| tagliaunghie | MO | NaN | 16 | 10981 | 35408.22 | 12081 | 19490.07 | 4 | 12 | 40.72 |
| tamburello | MO | 0 | 73 | 10981 | 60408 | 34588 | 35325.67 | 4 | 10 | 36.91 |
| tastiera | MO | 2 | 259 | 10981 | 97769.80 | 65653 | 58134.60 | 3 | 8 | 27 |
| telecomando | MO | 1 | 551 | 25753 | 69158.88 | 36447 | 43786.29 | 5 | 11 | 41.91 |
| temperino | MO | 1 | 40 | 25753 | 87376.17 | 133168 | 82099.06 | 4 | 9 | 32.43 |
| termometro | MO | 0 | 154 | 25753 | 71501 | 48641 | 48631.67 | 4 | 10 | 39.45 |
| trapano | MO | 2 | 254 | 29265 | 84975.25 | 133168 | 82469.42 | 3 | 7 | 26.33 |
| tubetto | MO | 2 | 96 | 16862 | 52790.75 | 223642 | 97764.92 | 3 | 7 | 25.48 |
| ventaglio | MO | 0 | 139 | 25262 | 108094.67 | 52265 | 61873.89 | 3 | 9 | 31.41 |
| volante | MO | 1 | 1900 | 15502 | 122185 | 86456 | 74714.33 | 3 | 7 | 25.4 |
| zuccheriera | MO | 1 | 14 | 333 | 85319.88 | 65653 | 50435.29 | 4 | 11 | 39.45 |
| angoscia | EMneg | 1 | 401 | 42551 | 41736 | 57191 | 47159.33 | 3 | 8 | 28.87 |
| antipatia | EMneg | 1 | 61 | 42551 | 98048.50 | 57191 | 65930.17 | 5 | 9 | 28.79 |
| avversione | EMneg | 1 | 104 | 32484 | 92905.29 | 70433 | 65274.10 | 4 | 10 | 35.48 |
| disagio | EMneg | 0 | 1521 | 174105 | 45165.25 | 52265 | 90511.75 | 3 | 7 | 23.03 |
| disgusto | EMneg | 1 | 231 | 174105 | 44772.60 | 223642 | 147506.53 | 3 | 8 | 28.62 |
| gelosia | EMneg | 1 | 782 | 8939 | 61503.50 | 57191 | 42544.50 | 4 | 7 | 23.96 |
| imbarazzo | EMneg | 2 | 1989 | 14207 | 60728.67 | 7038 | 27324.56 | 4 | 9 | 31.92 |
| infelicità | EMneg | 0 | 212 | 107539 | 41897 | 93005 | 80813.67 | 5 | 10 | 29.8 |
| inquietudine | EMneg | 1 | 53 | 107539 | 36583.67 | 70433 | 71518.56 | 5 | 12 | 39.45 |
| invidia | EMneg | 3 | 517 | 107539 | 41897 | 57191 | 68875.67 | 3 | 7 | 20.15 |
| malinconia | EMneg | 1 | 181 | 56674 | 91237.57 | 57191 | 68367.52 | 5 | 10 | 34.12 |
| nostalgia | EMneg | 1 | 566 | 52903 | 77439.33 | 57191 | 62511.11 | 4 | 9 | 31.41 |
| ostilità | EMneg | 0 | 368 | 3806 | 82699.40 | 93005 | 59836.80 | 4 | 8 | 24.81 |
| panico | EMneg | 2 | 2866 | 45016 | 107347 | 26810 | 59724.33 | 3 | 6 | 21 |
| rabbia | EMneg | 3 | 3431 | 16575 | 23626.67 | 57191 | 32464.22 | 2 | 6 | 20.07 |
| rammarico | EMneg | 1 | 191 | 16575 | 77057.17 | 26810 | 40147.39 | 4 | 9 | 33.7 |
| rancore | EMneg | 1 | 895 | 16575 | 104412 | 133974 | 84987 | 3 | 7 | 27.09 |
| repulsione | EMneg | 0 | 65 | 23590 | 55560.14 | 70433 | 49861.05 | 4 | 10 | 35.22 |
| rimorso | EMneg | 3 | 679 | 51082 | 57015.50 | 40259 | 49452.17 | 3 | 7 | 26.08 |
| rimpianto | EMneg | 2 | 459 | 51082 | 92641.33 | 223642 | 122455.11 | 3 | 9 | 30.56 |
| sdegno | EMneg | 1 | 90 | 170 | 34449.33 | 133168 | 55929.11 | 2 | 6 | 23.45 |
| sofferenza | EMneg | 1 | 1445 | 57963 | 77515.57 | 25784 | 53754.19 | 4 | 10 | 36.66 |
| terrore | EMneg | 2 | 1148 | 25753 | 108271.80 | 133974 | 89332.93 | 3 | 7 | 26.59 |
| tristezza | EMneg | 1 | 1109 | 29265 | 67533.17 | 25784 | 40860.72 | 3 | 9 | 30.06 |
| vergogna | EMneg | 3 | 2628 | 25262 | 53131.80 | 61022 | 46471.93 | 3 | 8 | 31.41 |
| affetto | EMpos | 6 | 1602 | 5695 | 56892.50 | 223642 | 95409.83 | 3 | 7 | 24.89 |
| allegria | EMpos | 1 | 371 | 68073 | 65022.40 | 57191 | 63428.80 | 4 | 8 | 26.84 |
| ammirazione | EMpos | 1 | 272 | 11832 | 74967.13 | 70433 | 52410.71 | 5 | 11 | 38.69 |
| divertimento | EMpos | 1 | 2717 | 174105 | 99486.56 | 223642 | 165744.52 | 5 | 12 | 41.57 |
| entusiasmo | EMpos | 2 | 853 | 4561 | 62046.86 | 27069 | 31225.62 | 4 | 10 | 36 |
| estasi | EMpos | 2 | 311 | 26683 | 98875 | 28994 | 51517.33 | 3 | 6 | 19.47 |
| euforia | EMpos | 1 | 128 | 1839 | 63701.25 | 57191 | 40910.42 | 4 | 7 | 24.3 |
| felicità | EMpos | 1 | 2929 | 8694 | 87007.20 | 93005 | 62902.07 | 4 | 8 | 24.13 |
| fiducia | EMpos | 0 | 5456 | 25684 | 26870.25 | 57191 | 36581.75 | 3 | 7 | 21.84 |
| gioia | EMpos | 4 | 3090 | 28311 | 35936 | 57191 | 40479.33 | 2 | 5 | 15.66 |
| gradimento | EMpos | 1 | 257 | 14632 | 93770.29 | 223642 | 110681.43 | 4 | 10 | 37 |
| gratitudine | EMpos | 1 | 775 | 14632 | 86536.13 | 70433 | 57200.38 | 5 | 11 | 37.25 |
| ilarità | EMpos | 0 | 20 | 67973 | 98161.75 | 93005 | 86379.92 | 4 | 7 | 21 |
| letizia | EMpos | 0 | 62 | 38814 | 57952.75 | 57191 | 51319.25 | 3 | 7 | 20.48 |
| meraviglia | EMpos | 3 | 1939 | 28080 | 73575.57 | 57191 | 52948.86 | 4 | 10 | 33.6 |
| ottimismo | EMpos | 0 | 200 | 2888 | 61817.17 | 27069 | 30591.39 | 4 | 9 | 31.16 |
| passione | EMpos | 1 | 2456 | 45016 | 99446 | 70433 | 71631.67 | 3 | 8 | 28.11 |
| riconoscenza | EMpos | 1 | 163 | 51082 | 82940.33 | 25784 | 53268.78 | 5 | 12 | 44 |
| serenità | EMpos | 0 | 246 | 58313 | 123363 | 93005 | 91560.33 | 4 | 8 | 27.52 |
| simpatia | EMpos | 1 | 440 | 50691 | 79781 | 57191 | 62554.33 | 4 | 8 | 25.57 |
| soddisfazione | EMpos | 1 | 805 | 57963 | 54687.30 | 70433 | 61027.77 | 5 | 13 | 44.79 |
| solievo | EMpos | 0 | 1189 | 57963 | 77574 | 12497 | 49344.67 | 3 | 8 | 27.09 |
| stima | EMpos | 5 | 742 | 49172 | 63435.50 | 22625 | 45077.50 | 2 | 5 | 16.93 |
| tenerezza | EMpos | 1 | 271 | 25753 | 88882 | 25784 | 46806.33 | 4 | 9 | 32.43 |
| tranquillità | EMpos | 0 | 454 | 29265 | 76285.11 | 93005 | 66185.04 | 4 | 12 | 39.54 |
| abitudine | NEU | 1 | 1455 | 9465 | 51751.50 | 70433 | 43883.17 | 5 | 9 | 29.38 |
| buonsenso | NEU | 0 | 187 | 4458 | 82890.33 | 40259 | 42535.78 | 3 | 9 | 34.71 |
| cautela | NEU | 1 | 426 | 46052 | 54334.50 | 78467 | 59617.83 | 3 | 7 | 24.98 |
| contegno | NEU | 2 | 135 | 135447 | 104467 | 133168 | 124360.67 | 3 | 8 | 31.41 |
| criterio | NEU | 0 | 117 | 9576 | 115488.80 | 52265 | 59109.93 | 3 | 8 | 26.33 |
| decenza | NEU | 3 | 269 | 116530 | 70689.25 | 25784 | 71001.08 | 3 | 7 | 25.99 |
| discrezione | NEU | 0 | 656 | 174105 | 73872.25 | 70433 | 106136.75 | 4 | 11 | 37.51 |
| giudizio | NEU | 0 | 2352 | 28311 | 110561 | 52265 | 63712.33 | 3 | 8 | 24.55 |
| imparzialità | NEU | 0 | 53 | 14207 | 71884.78 | 93005 | 59698.93 | 5 | 12 | 37.76 |
| impressione | NEU | 2 | 2782 | 14207 | 89438.50 | 70433 | 58026.17 | 4 | 11 | 38.02 |
| memoria | NEU | 2 | 5175 | 28080 | 74617.50 | 57191 | 53296.17 | 3 | 7 | 26 |
| moderazione | NEU | 0 | 106 | 29591 | 96005.88 | 70433 | 65343.29 | 5 | 11 | 40.47 |
| neutralità | NEU | 0 | 34 | 40713 | 76388.43 | 93005 | 70035.48 | 4 | 10 | 34.21 |
| normalità | NEU | 1 | 620 | 52903 | 78091.17 | 93005 | 74666.39 | 4 | 9 | 32.09 |
| obiettività | NEU | 0 | 66 | 1240 | 64806.50 | 93005 | 53017.17 | 5 | 11 | 33.95 |
| opinione | NEU | 1 | 3404 | 4461 | 94509 | 70433 | 56467.67 | 4 | 8 | 26.8 |
| parsimonia | NEU | 0 | 24 | 45016 | 78901.43 | 57191 | 60369.48 | 4 | 10 | 34.46 |
| pensiero | NEU | 1 | 4320 | 74312 | 102951 | 48641 | 75301.33 | 3 | 8 | 28.28 |
| presentimento | NEU | 1 | 785 | 66996 | 115006.20 | 223642 | 135214.73 | 5 | 13 | 47.07 |
| prudenza | NEU | 1 | 520 | 66996 | 53890.60 | 25784 | 48890.20 | 3 | 8 | 29.89 |
| quotidianità | NEU | 0 | 21 | 59294 | 68089.22 | 93005 | 73462.74 | 5 | 12 | 39.2 |
| raziocinio | NEU | 0 | 21 | 16575 | 62232.60 | 52265 | 43690.87 | 4 | 10 | 32 |
| regolarità | NEU | 0 | 89 | 23590 | 77927.71 | 93005 | 64840.90 | 5 | 10 | 35.22 |
| senno | NEU | 7 | 416 | 58313 | 113335 | 133168 | 101605.33 | 2 | 5 | 19 |
| sobrietà | NEU | 0 | 125 | 57963 | 47763.20 | 93005 | 66243.73 | 3 | 8 | 27.77 |

**Supplementary Table 1B.**
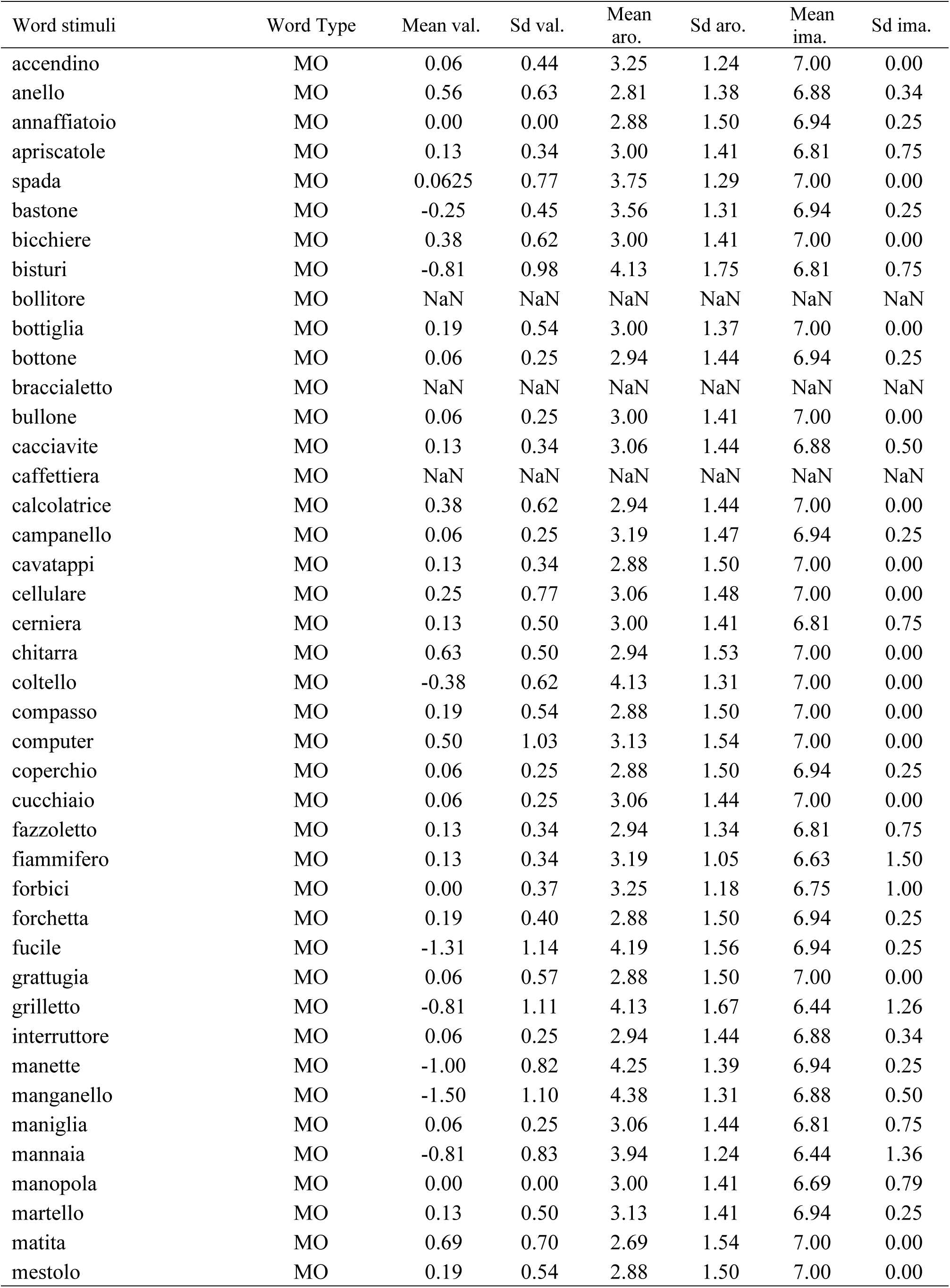
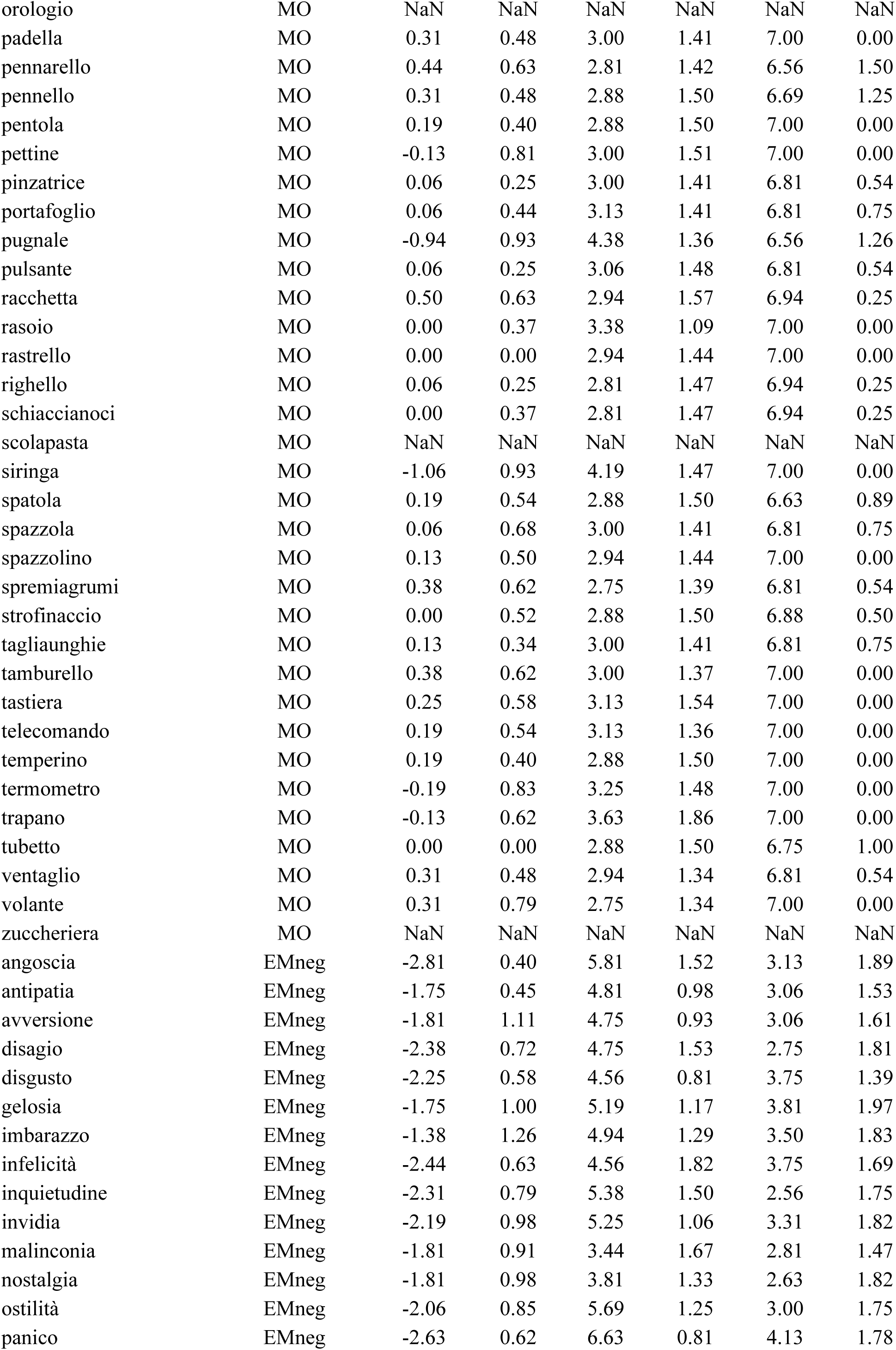
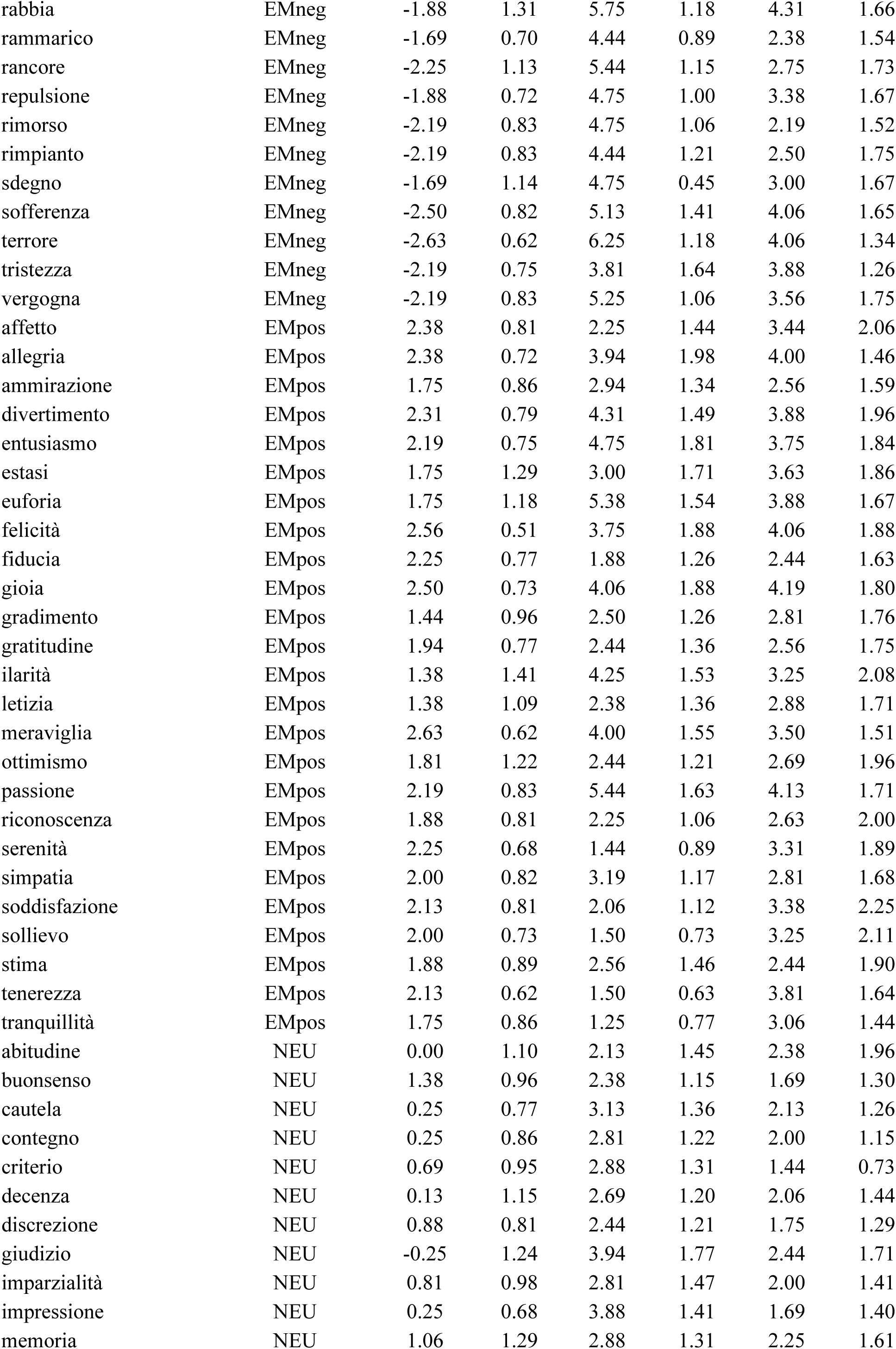
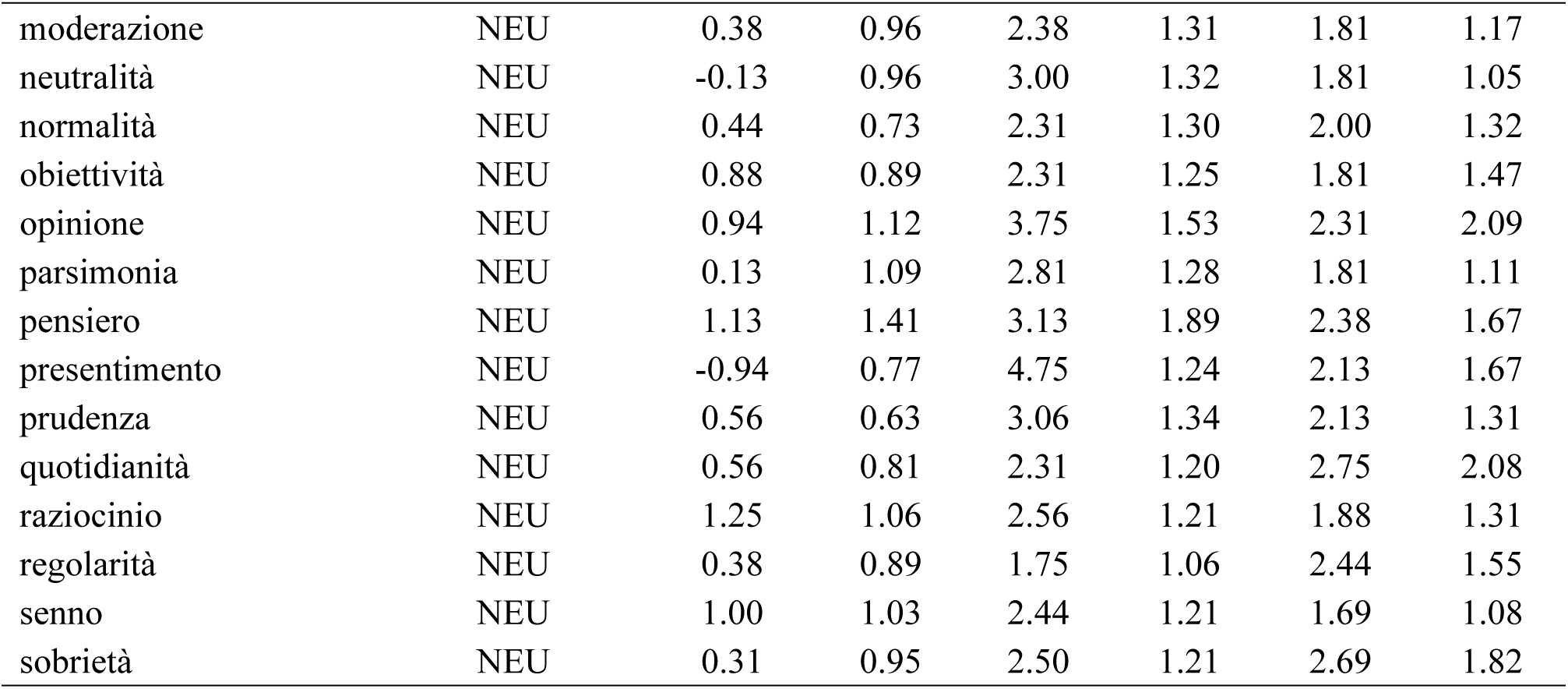
Descriptive statistics of the rating scores for valence (val.), arousal (aro.), and imaginability (ima.) for every experimental stimulus obtained in the preliminary rating study on independent sample of 16 participants.

| Word stimuli | Word Type | Mean val. | Sd val. | Mean aro. | Sd aro. | Mean ima. | Sd ima. |
| --- | --- | --- | --- | --- | --- | --- | --- |
| accendino | MO | 0.06 | 0.44 | 3.25 | 1.24 | 7.00 | 0.00 |
| anello | MO | 0.56 | 0.63 | 2.81 | 1.38 | 6.88 | 0.34 |
| annaffiatoio | MO | 0.00 | 0.00 | 2.88 | 1.50 | 6.94 | 0.25 |
| apricatole | MO | 0.13 | 0.34 | 3.00 | 1.41 | 6.81 | 0.75 |
| spada | MO | 0.0625 | 0.77 | 3.75 | 1.29 | 7.00 | 0.00 |
| bastone | MO | -0.25 | 0.45 | 3.56 | 1.31 | 6.94 | 0.25 |
| bicchiera | MO | 0.38 | 0.62 | 3.00 | 1.41 | 7.00 | 0.00 |
| bisturi | MO | -0.81 | 0.98 | 4.13 | 1.75 | 6.81 | 0.75 |
| bollitore | MO | NaN | NaN | NaN | NaN | NaN | NaN |
| bottiglia | MO | 0.19 | 0.54 | 3.00 | 1.37 | 7.00 | 0.00 |
| bottone | MO | 0.06 | 0.25 | 2.94 | 1.44 | 6.94 | 0.25 |
| braccialeto | MO | NaN | NaN | NaN | NaN | NaN | NaN |
| bullone | MO | 0.06 | 0.25 | 3.00 | 1.41 | 7.00 | 0.00 |
| cacciavite | MO | 0.13 | 0.34 | 3.06 | 1.44 | 6.88 | 0.50 |
| caffettiera | MO | NaN | NaN | NaN | NaN | NaN | NaN |
| calcolatrice | MO | 0.38 | 0.62 | 2.94 | 1.44 | 7.00 | 0.00 |
| campanello | MO | 0.06 | 0.25 | 3.19 | 1.47 | 6.94 | 0.25 |
| cavatappi | MO | 0.13 | 0.34 | 2.88 | 1.50 | 7.00 | 0.00 |
| cellulare | MO | 0.25 | 0.77 | 3.06 | 1.48 | 7.00 | 0.00 |
| cerniera | MO | 0.13 | 0.50 | 3.00 | 1.41 | 6.81 | 0.75 |
| chitarra | MO | 0.63 | 0.50 | 2.94 | 1.53 | 7.00 | 0.00 |
| coltello | MO | -0.38 | 0.62 | 4.13 | 1.31 | 7.00 | 0.00 |
| compasso | MO | 0.19 | 0.54 | 2.88 | 1.50 | 7.00 | 0.00 |
| computer | MO | 0.50 | 1.03 | 3.13 | 1.54 | 7.00 | 0.00 |
| coperchio | MO | 0.06 | 0.25 | 2.88 | 1.50 | 6.94 | 0.25 |
| cucchiaio | MO | 0.06 | 0.25 | 3.06 | 1.44 | 7.00 | 0.00 |
| fazzoletto | MO | 0.13 | 0.34 | 2.94 | 1.34 | 6.81 | 0.75 |
| fiammifero | MO | 0.13 | 0.34 | 3.19 | 1.05 | 6.63 | 1.50 |
| forbici | MO | 0.00 | 0.37 | 3.25 | 1.18 | 6.75 | 1.00 |
| forchetta | MO | 0.19 | 0.40 | 2.88 | 1.50 | 6.94 | 0.25 |
| fucile | MO | -1.31 | 1.14 | 4.19 | 1.56 | 6.94 | 0.25 |
| grattugia | MO | 0.06 | 0.57 | 2.88 | 1.50 | 7.00 | 0.00 |
| grilletto | MO | -0.81 | 1.11 | 4.13 | 1.67 | 6.44 | 1.26 |
| interruttore | MO | 0.06 | 0.25 | 2.94 | 1.44 | 6.88 | 0.34 |
| manette | MO | -1.00 | 0.82 | 4.25 | 1.39 | 6.94 | 0.25 |
| manganello | MO | -1.50 | 1.10 | 4.38 | 1.31 | 6.88 | 0.50 |
| maniglia | MO | 0.06 | 0.25 | 3.06 | 1.44 | 6.81 | 0.75 |
| mannaia | MO | -0.81 | 0.83 | 3.94 | 1.24 | 6.44 | 1.36 |
| manopola | MO | 0.00 | 0.00 | 3.00 | 1.41 | 6.69 | 0.79 |
| martello | MO | 0.13 | 0.50 | 3.13 | 1.41 | 6.94 | 0.25 |
| matita | MO | 0.69 | 0.70 | 2.69 | 1.54 | 7.00 | 0.00 |
| mestolo | MO | 0.19 | 0.54 | 2.88 | 1.50 | 7.00 | 0.00 |
| orologio | MO | NaN | NaN | NaN | NaN | NaN | NaN |
| padella | MO | 0.31 | 0.48 | 3.00 | 1.41 | 7.00 | 0.00 |
| pennarello | MO | 0.44 | 0.63 | 2.81 | 1.42 | 6.56 | 1.50 |
| pennello | MO | 0.31 | 0.48 | 2.88 | 1.50 | 6.69 | 1.25 |
| pentola | MO | 0.19 | 0.40 | 2.88 | 1.50 | 7.00 | 0.00 |
| pettine | MO | -0.13 | 0.81 | 3.00 | 1.51 | 7.00 | 0.00 |
| pinzatrice | MO | 0.06 | 0.25 | 3.00 | 1.41 | 6.81 | 0.54 |
| portafoglio | MO | 0.06 | 0.44 | 3.13 | 1.41 | 6.81 | 0.75 |
| pugnale | MO | -0.94 | 0.93 | 4.38 | 1.36 | 6.56 | 1.26 |
| pulsante | MO | 0.06 | 0.25 | 3.06 | 1.48 | 6.81 | 0.54 |
| racchetta | MO | 0.50 | 0.63 | 2.94 | 1.57 | 6.94 | 0.25 |
| rasoio | MO | 0.00 | 0.37 | 3.38 | 1.09 | 7.00 | 0.00 |
| rastrello | MO | 0.00 | 0.00 | 2.94 | 1.44 | 7.00 | 0.00 |
| righello | MO | 0.06 | 0.25 | 2.81 | 1.47 | 6.94 | 0.25 |
| schiaccianoci | MO | 0.00 | 0.37 | 2.81 | 1.47 | 6.94 | 0.25 |
| scolapasta | MO | NaN | NaN | NaN | NaN | NaN | NaN |
| siringa | MO | -1.06 | 0.93 | 4.19 | 1.47 | 7.00 | 0.00 |
| spatola | MO | 0.19 | 0.54 | 2.88 | 1.50 | 6.63 | 0.89 |
| spazzola | MO | 0.06 | 0.68 | 3.00 | 1.41 | 6.81 | 0.75 |
| spazzolino | MO | 0.13 | 0.50 | 2.94 | 1.44 | 7.00 | 0.00 |
| spremiagrumi | MO | 0.38 | 0.62 | 2.75 | 1.39 | 6.81 | 0.54 |
| strofinaccio | MO | 0.00 | 0.52 | 2.88 | 1.50 | 6.88 | 0.50 |
| tagliaunghie | MO | 0.13 | 0.34 | 3.00 | 1.41 | 6.81 | 0.75 |
| tamburello | MO | 0.38 | 0.62 | 3.00 | 1.37 | 7.00 | 0.00 |
| tastiera | MO | 0.25 | 0.58 | 3.13 | 1.54 | 7.00 | 0.00 |
| telecomando | MO | 0.19 | 0.54 | 3.13 | 1.36 | 7.00 | 0.00 |
| temperino | MO | 0.19 | 0.40 | 2.88 | 1.50 | 7.00 | 0.00 |
| termometro | MO | -0.19 | 0.83 | 3.25 | 1.48 | 7.00 | 0.00 |
| trapano | MO | -0.13 | 0.62 | 3.63 | 1.86 | 7.00 | 0.00 |
| tubetto | MO | 0.00 | 0.00 | 2.88 | 1.50 | 6.75 | 1.00 |
| ventaglio | MO | 0.31 | 0.48 | 2.94 | 1.34 | 6.81 | 0.54 |
| volante | MO | 0.31 | 0.79 | 2.75 | 1.34 | 7.00 | 0.00 |
| zuccheriera | MO | NaN | NaN | NaN | NaN | NaN | NaN |
| angoscia | EMneg | -2.81 | 0.40 | 5.81 | 1.52 | 3.13 | 1.89 |
| antipatia | EMneg | -1.75 | 0.45 | 4.81 | 0.98 | 3.06 | 1.53 |
| avversione | EMneg | -1.81 | 1.11 | 4.75 | 0.93 | 3.06 | 1.61 |
| disagio | EMneg | -2.38 | 0.72 | 4.75 | 1.53 | 2.75 | 1.81 |
| disgusto | EMneg | -2.25 | 0.58 | 4.56 | 0.81 | 3.75 | 1.39 |
| gelosia | EMneg | -1.75 | 1.00 | 5.19 | 1.17 | 3.81 | 1.97 |
| imbarazzo | EMneg | -1.38 | 1.26 | 4.94 | 1.29 | 3.50 | 1.83 |
| infelicità | EMneg | -2.44 | 0.63 | 4.56 | 1.82 | 3.75 | 1.69 |
| inquietudine | EMneg | -2.31 | 0.79 | 5.38 | 1.50 | 2.56 | 1.75 |
| invidia | EMneg | -2.19 | 0.98 | 5.25 | 1.06 | 3.31 | 1.82 |
| malinconia | EMneg | -1.81 | 0.91 | 3.44 | 1.67 | 2.81 | 1.47 |
| nostalgia | EMneg | -1.81 | 0.98 | 3.81 | 1.33 | 2.63 | 1.82 |
| ostilità | EMneg | -2.06 | 0.85 | 5.69 | 1.25 | 3.00 | 1.75 |
| panico | EMneg | -2.63 | 0.62 | 6.63 | 0.81 | 4.13 | 1.78 |
| rabbia | EMneg | -1.88 | 1.31 | 5.75 | 1.18 | 4.31 | 1.66 |
| rammarico | EMneg | -1.69 | 0.70 | 4.44 | 0.89 | 2.38 | 1.54 |
| rancore | EMneg | -2.25 | 1.13 | 5.44 | 1.15 | 2.75 | 1.73 |
| repulsione | EMneg | -1.88 | 0.72 | 4.75 | 1.00 | 3.38 | 1.67 |
| rimorso | EMneg | -2.19 | 0.83 | 4.75 | 1.06 | 2.19 | 1.52 |
| rimpianto | EMneg | -2.19 | 0.83 | 4.44 | 1.21 | 2.50 | 1.75 |
| sdegno | EMneg | -1.69 | 1.14 | 4.75 | 0.45 | 3.00 | 1.67 |
| sofferenza | EMneg | -2.50 | 0.82 | 5.13 | 1.41 | 4.06 | 1.65 |
| terrore | EMneg | -2.63 | 0.62 | 6.25 | 1.18 | 4.06 | 1.34 |
| tristezza | EMneg | -2.19 | 0.75 | 3.81 | 1.64 | 3.88 | 1.26 |
| vergogna | EMneg | -2.19 | 0.83 | 5.25 | 1.06 | 3.56 | 1.75 |
| affetto | EMpos | 2.38 | 0.81 | 2.25 | 1.44 | 3.44 | 2.06 |
| allegria | EMpos | 2.38 | 0.72 | 3.94 | 1.98 | 4.00 | 1.46 |
| ammirazione | EMpos | 1.75 | 0.86 | 2.94 | 1.34 | 2.56 | 1.59 |
| divertimento | EMpos | 2.31 | 0.79 | 4.31 | 1.49 | 3.88 | 1.96 |
| entusiasmo | EMpos | 2.19 | 0.75 | 4.75 | 1.81 | 3.75 | 1.84 |
| estasi | EMpos | 1.75 | 1.29 | 3.00 | 1.71 | 3.63 | 1.86 |
| euforia | EMpos | 1.75 | 1.18 | 5.38 | 1.54 | 3.88 | 1.67 |
| felicità | EMpos | 2.56 | 0.51 | 3.75 | 1.88 | 4.06 | 1.88 |
| fiducia | EMpos | 2.25 | 0.77 | 1.88 | 1.26 | 2.44 | 1.63 |
| gioia | EMpos | 2.50 | 0.73 | 4.06 | 1.88 | 4.19 | 1.80 |
| gradimento | EMpos | 1.44 | 0.96 | 2.50 | 1.26 | 2.81 | 1.76 |
| gratitudine | EMpos | 1.94 | 0.77 | 2.44 | 1.36 | 2.56 | 1.75 |
| ilarità | EMpos | 1.38 | 1.41 | 4.25 | 1.53 | 3.25 | 2.08 |
| letizia | EMpos | 1.38 | 1.09 | 2.38 | 1.36 | 2.88 | 1.71 |
| meraviglia | EMpos | 2.63 | 0.62 | 4.00 | 1.55 | 3.50 | 1.51 |
| ottimismo | EMpos | 1.81 | 1.22 | 2.44 | 1.21 | 2.69 | 1.96 |
| passione | EMpos | 2.19 | 0.83 | 5.44 | 1.63 | 4.13 | 1.71 |
| riconoscenza | EMpos | 1.88 | 0.81 | 2.25 | 1.06 | 2.63 | 2.00 |
| serenità | EMpos | 2.25 | 0.68 | 1.44 | 0.89 | 3.31 | 1.89 |
| simpatia | EMpos | 2.00 | 0.82 | 3.19 | 1.17 | 2.81 | 1.68 |
| soddisfazione | EMpos | 2.13 | 0.81 | 2.06 | 1.12 | 3.38 | 2.25 |
| solievo | EMpos | 2.00 | 0.73 | 1.50 | 0.73 | 3.25 | 2.11 |
| stima | EMpos | 1.88 | 0.89 | 2.56 | 1.46 | 2.44 | 1.90 |
| tenerezza | EMpos | 2.13 | 0.62 | 1.50 | 0.63 | 3.81 | 1.64 |
| tranquillità | EMpos | 1.75 | 0.86 | 1.25 | 0.77 | 3.06 | 1.44 |
| abitudine | NEU | 0.00 | 1.10 | 2.13 | 1.45 | 2.38 | 1.96 |
| buonsenso | NEU | 1.38 | 0.96 | 2.38 | 1.15 | 1.69 | 1.30 |
| cautela | NEU | 0.25 | 0.77 | 3.13 | 1.36 | 2.13 | 1.26 |
| contegno | NEU | 0.25 | 0.86 | 2.81 | 1.22 | 2.00 | 1.15 |
| criterio | NEU | 0.69 | 0.95 | 2.88 | 1.31 | 1.44 | 0.73 |
| decenza | NEU | 0.13 | 1.15 | 2.69 | 1.20 | 2.06 | 1.44 |
| discrezione | NEU | 0.88 | 0.81 | 2.44 | 1.21 | 1.75 | 1.29 |
| giudizio | NEU | -0.25 | 1.24 | 3.94 | 1.77 | 2.44 | 1.71 |
| imparzialità | NEU | 0.81 | 0.98 | 2.81 | 1.47 | 2.00 | 1.41 |
| impressione | NEU | 0.25 | 0.68 | 3.88 | 1.41 | 1.69 | 1.40 |
| memoria | NEU | 1.06 | 1.29 | 2.88 | 1.31 | 2.25 | 1.61 |
| moderazione | NEU | 0.38 | 0.96 | 2.38 | 1.31 | 1.81 | 1.17 |
| neutralità | NEU | -0.13 | 0.96 | 3.00 | 1.32 | 1.81 | 1.05 |
| normalità | NEU | 0.44 | 0.73 | 2.31 | 1.30 | 2.00 | 1.32 |
| obiettività | NEU | 0.88 | 0.89 | 2.31 | 1.25 | 1.81 | 1.47 |
| opinione | NEU | 0.94 | 1.12 | 3.75 | 1.53 | 2.31 | 2.09 |
| parsimonia | NEU | 0.13 | 1.09 | 2.81 | 1.28 | 1.81 | 1.11 |
| pensiero | NEU | 1.13 | 1.41 | 3.13 | 1.89 | 2.38 | 1.67 |
| presentimento | NEU | -0.94 | 0.77 | 4.75 | 1.24 | 2.13 | 1.67 |
| prudenza | NEU | 0.56 | 0.63 | 3.06 | 1.34 | 2.13 | 1.31 |
| quotidianità | NEU | 0.56 | 0.81 | 2.31 | 1.20 | 2.75 | 2.08 |
| raziocinio | NEU | 1.25 | 1.06 | 2.56 | 1.21 | 1.88 | 1.31 |
| regolarità | NEU | 0.38 | 0.89 | 1.75 | 1.06 | 2.44 | 1.55 |
| senno | NEU | 1.00 | 1.03 | 2.44 | 1.21 | 1.69 | 1.08 |
| sobrietà | NEU | 0.31 | 0.95 | 2.50 | 1.21 | 2.69 | 1.82 |

**Supplementary Table 2A.**
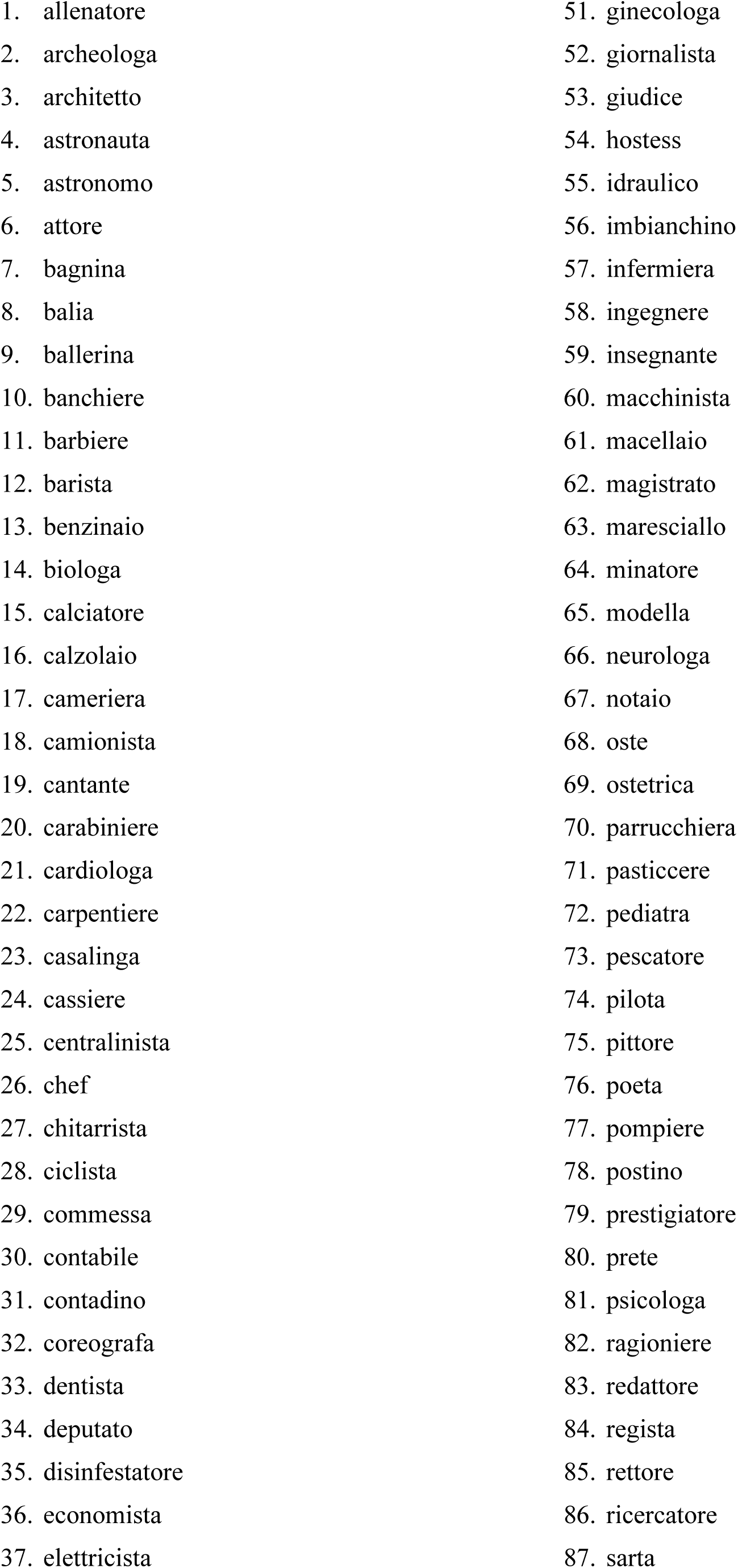
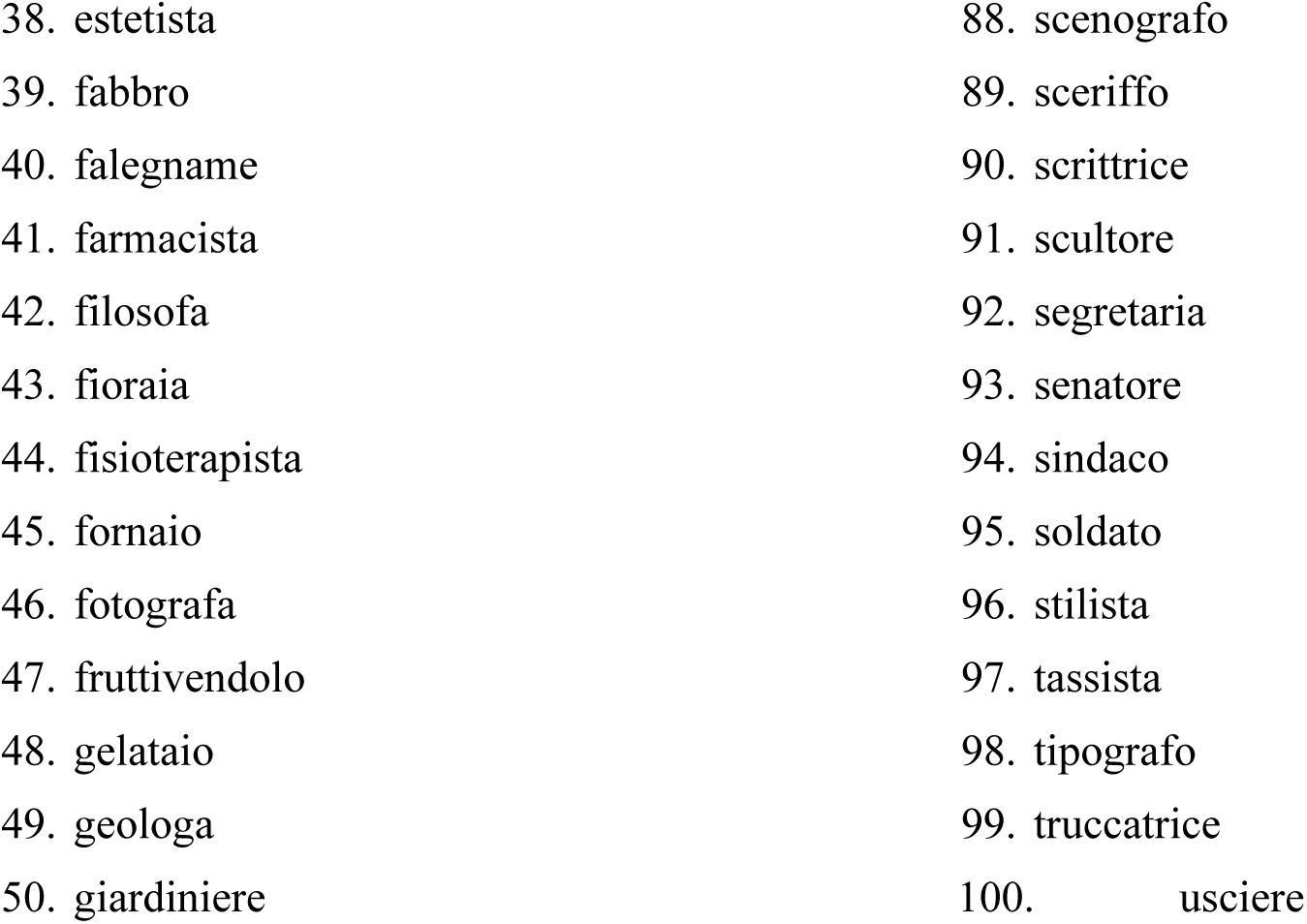
100 words referring to jobs and careers used in the training session.

|  |  |
| --- | --- |
| 1. allenatore | 51. ginecologa |
| 2. archeologa | 52. giornalista |
| 3. architetto | 53. giudice |
| 4. astronauta | 54. hostess |
| 5. astronomo | 55. idraulico |
| 6. attore | 56. imbianchino |
| 7. bagnina | 57. infermiera |
| 8. balia | 58. ingegnere |
| 9. ballerina | 59. insegnante |
| 10. banchiere | 60. macchinista |
| 11. barbiere | 61. macellaio |
| 12. barista | 62. magistrato |
| 13. benzinaio | 63. maresciallo |
| 14. biologa | 64. minatore |
| 15. calciatore | 65. modella |
| 16. calzolaio | 66. neurologa |
| 17. cameriera | 67. notaio |
| 18. camionista | 68. oste |
| 19. cantante | 69. ostetrica |
| 20. carabiniere | 70. parrucchiera |
| 21. cardiologa | 71. pasticcere |
| 22. carpentiere | 72. pediatra |
| 23. casalinga | 73. pescatore |
| 24. cassiere | 74. pilota |
| 25. centralinista | 75. pittore |
| 26. chef | 76. poeta |
| 27. chitarrista | 77. pompiere |
| 28. ciclista | 78. postino |
| 29. commessa | 79. prestigiatore |
| 30. contabile | 80. prete |
| 31. contadino | 81. psicologa |
| 32. coreografa | 82. ragioniere |
| 33. dentista | 83. redattore |
| 34. deputato | 84. regista |
| 35. disinfestatore | 85. rettore |
| 36. economista | 86. ricercatore |
| 37. elettricista | 87. sarta |
| 38. estetista | 88. scenografo |
| 39. fabbro | 89. sceriffo |
| 40. falegname | 90. scrittrice |
| 41. farmacista | 91. scultore |
| 42. filosofa | 92. segretaria |
| 43. fioraia | 93. senatore |
| 44. fisioterapista | 94. sindaco |
| 45. fornaio | 95. soldato |
| 46. fotografa | 96. stilista |
| 47. fruttivendolo | 97. tassista |
| 48. gelataio | 98. tipografo |
| 49. geologa | 99. truccatrice |
| 50. giardiniere | 100. usciera |

**Supplementary Table 2B.** Lexical and sublexical variables of training stimuli referring to jobs and careers, matched with manipulable objects (MO) and emotion (EM, with negatively valenced [EMneg], positively valenced [EMpos], and neutral [NEU]) word stimuli, mean (sd).

| Variable | MO | EMneg | EMpos | NEU | Career | Kruskal-Wallis | *p* value |
| --- | --- | --- | --- | --- | --- | --- | --- |
| Lexical frequency | 975.4<br>(1663) | 879.7<br>(949.1) | 1109.9<br>(1319.3) | 966.3<br>(1462.2) | 770.43<br>(1254.557) | 5.6171 | 0.2296 |
| N. syllables | 3.48<br>(0.67) | 3.56<br>(0.87) | 3.8<br>(0.87) | 3.8<br>(0.91) | 3.69<br>(0.87) | 6.8592 | 0.143 |
| N. letters | 8.76<br>(1.8) | 8.32<br>(1.55) | 8.72<br>(2.23) | 9.2<br>(1.91) | 8.8<br>(2.01) | 2.7892 | 0.56 |

**Supplementary Table 3A.**
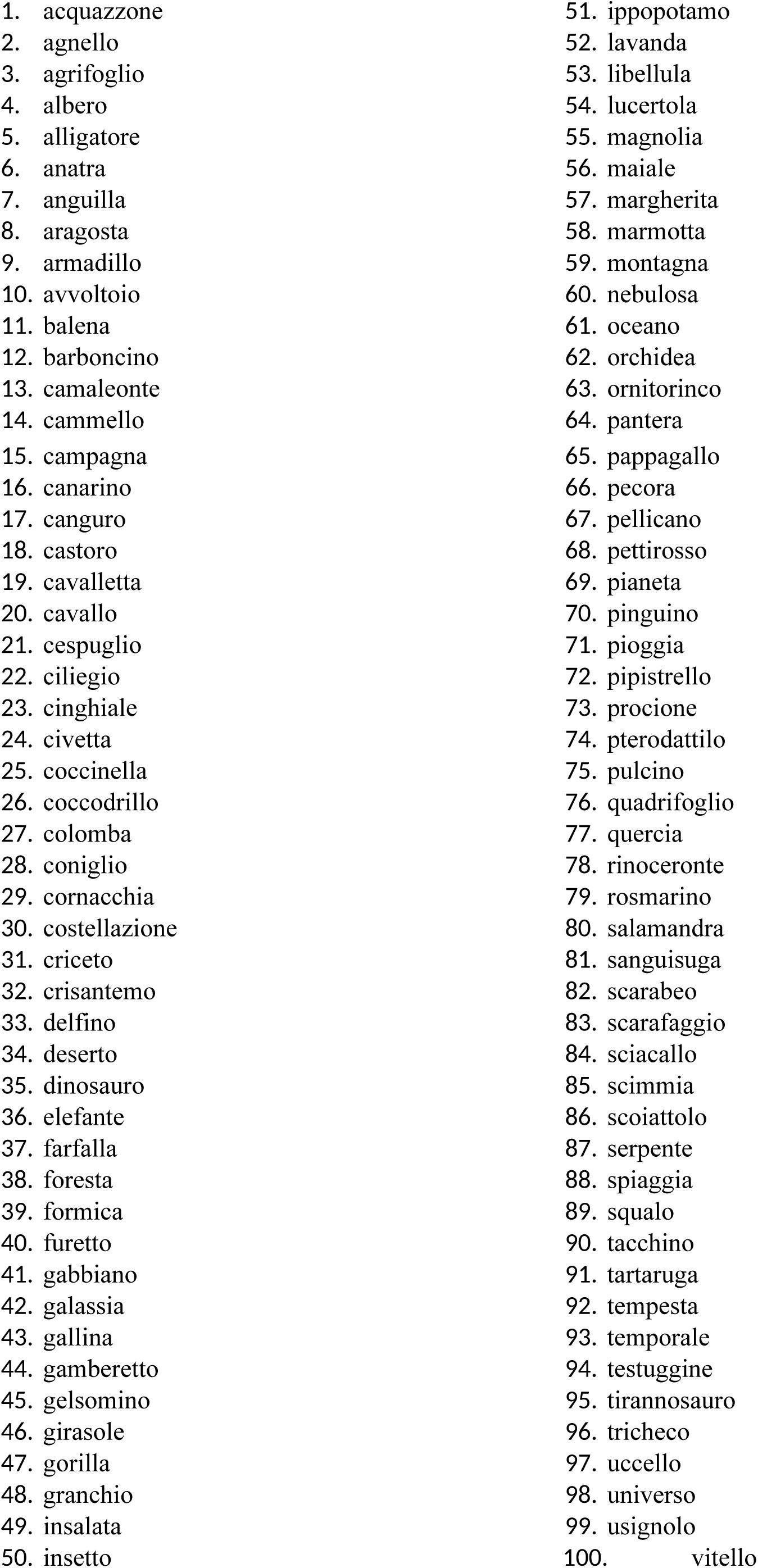
100 words referring to living/natural entities used to assess participants’ perceptual threshold.

**Supplementary Table 3B.** Lexical and sublexical and psycholinguistic variables of the stimuli referring living/natural items used for assessing the participants’ perceptual threshold, matched with manipulable objects (MO) and emotion (EM, with negatively valenced [EMneg], positively valenced [EMpos], and neutral [NEU]) word stimuli, mean (sd).

| Variable | MO | EMneg | EMpos | NEU | Living | Kruskal-Wallis | *p* value |
| --- | --- | --- | --- | --- | --- | --- | --- |
| Lexical frequency | 975.4<br>(1663) | 879.7<br>(949.1) | 1109.9<br>(1319.3) | 966.3<br>(1462.2) | 844.74<br>(1372) | 5.2586 | 0.262 |
| First bigram frequency | 36399.6<br>(32386.9) | 51354<br>(47659.77) | 37908.56<br>(35402.45) | 47136<br>(42814.64) | 40578.68<br>(38356.7) | 2.3387 | 0.6737 |
| Last bigram frequency | 70599.56<br>(49678.53) | 76472.72<br>(55314.3) | 75793.48<br>(60578.25) | 79331.6<br>(40676.11) | 72701.65<br>(48623.17) | 2.3375 | 0.674 |
| Medial bigrams frequency | 74687.16<br>(22995.85) | 67007<br>(25336.21) | 75560.52<br>(21534.27) | 81195.73<br>(20707.4) | 71242.46<br>(23138.97) | 6.4627 | 0.1672 |
| Mean bigram frequency | 60562.12<br>(20598.21) | 64944.58<br>(27122.55) | 63087.52<br>(29200.52) | 69221.12<br>(23766.04) | 61507.6<br>(23470.87) | 2.6986 | 0.601 |
| N. orthographic neighbours | 1.51<br>(1.51) | 1.32<br>(0.94) | 1.36<br>(1.58) | 0.92<br>(1.52) | 1.4<br>(1.188) | 7.7002 | 0.103 |
| N. syllables | 3.48<br>(0.67) | 3.56<br>(0.87) | 3.8<br>(0.87) | 3.8<br>(0.91) | 3.5<br>(0.76) | 6.2544 | 0.181 |
| N. letters | 8.76<br>(1.8) | 8.32<br>(1.55) | 8.72<br>(2.23) | 9.2<br>(1.91) | 8.43<br>(1.55) | 4.7944 | 0.309 |

